# Structure-constrained spin responses of an FMN–superoxide radical-pair model in mitochondrial complex I

**DOI:** 10.64898/2026.07.15.738821

**Authors:** Ji-Yong Sung, Lewis M. Antill, Jae-Ho Cheong

## Abstract

Mitochondrial complex I produces reactive oxygen species at its flavin mononucleotide (FMN) cofactor. We combined steric sampling of oxygen positions in human complex I with radical-pair spin simulations to examine how the FMN pocket shapes reaction-yield sensitivity, which provided singlet yields across FMN–O_2_ centroid separations, magnetic fields and spin-dephasing conditions. When candidate oxygen positions overlapping the FMN cofactor itself were excluded, almost no accessible positions remained within 3.3 Å of the isoalloxazine centre (≤0.2%), and a composite score combining geometric accessibility with normalised field and dephasing responses had no near-contact maximum. In the reference structure (PDB 5XTD), the highest-scoring 10% of accessible configurations had a mean separation of 5.8 Å (2.5th–97.5th percentiles, 5.6–6.0 Å); in the independently determined 9TI4 structure, the corresponding mean was 7.9 Å (7.7–8.0 Å), reflecting differences in the modelled pocket. Within each structure, this location changed little when the dephasing model, flavin hyperfine parameters, steric radii and recombination/dephasing rates were varied. Over 2.5–20 Å, the exchange-only score formed a broad plateau without a localized maximum. Small prescribed fluctuations in separation produced less singlet-yield variability in the high-scoring configurations than elsewhere in the accessible ensemble. A near-contact score maximum reported in an earlier version of this analysis arose because positions overlapping FMN had been counted as accessible. The accessibility-weighted spin-response map of the complex I flavin site is therefore set mainly by pocket geometry and should be evaluated over structural ensembles rather than a single model.

## Introduction

Mitochondrial reactive oxygen species (ROS) participate in cell signalling, metabolic responses and oxidative stress.(1, 2) The respiratory-chain enzyme complex I is a major source of mitochondrial superoxide, with the flavin mononucleotide (FMN) cofactor among the sites where molecular oxygen undergoes reduction.(3, 13, 14) Electron-transfer kinetics, redox state and oxygen access influence ROS formation at this site. The possibility that radical-pair spin dynamics also affect the distribution of reaction products can be studied by combining the local protein geometry with calculations of singlet–triplet evolution.(7, 9, 18)

Complex I transfers electrons from NADH through FMN and iron–sulfur centres to ubiquinone and contributes to proton translocation.(13) Transfer of an electron from reduced FMN to triplet-ground-state oxygen can generate a spin-correlated FMN semiquinone– superoxide radical pair.(14, 15, 33) In a proposed reaction scheme, the triplet channel releases superoxide, whereas an additional electron transfer through the singlet channel produces hydrogen peroxide.(33) The relative yields may depend on hyperfine coupling, electron– electron interactions and spin relaxation during the radical-pair lifetime.

Molecular-dynamics simulations have located oxygen near flavin cofactors in electron-transfer flavoprotein and *A. thaliana* cryptochrome 1.(42, 43) The cryptochrome calculations also describe persistent superoxide configurations.(43) Such observations provide a basis for studying how the shape of a flavin pocket selects oxygen-approach geometries and may influence the motion of a radical formed after electron transfer.

The magnetic response of a superoxide-containing radical pair also depends on the motion of the superoxide radical. Spin–orbit coupling makes its electron-spin relaxation sensitive to molecular reorientation.(40) Simulations of avian cryptochrome 4 show that association with a protein can slow superoxide rotation.(41) This motivates examining a range of spin-dephasing rates when modelling radical-pair reactions in complex I.

The FMN pocket restricts the separations and orientations from which oxygen can approach the cofactor. Exchange coupling changes rapidly with radical separation, dipolar coupling depends on separation and orientation, and flavin hyperfine interactions provide an additional source of singlet–triplet evolution.(16, 34) Mapping accessible oxygen positions and the associated spin-response profiles therefore gives a spatial account of the configurations available for radical-pair chemistry.

We sampled oxygen positions around FMN from the human complex I cryo-EM structures 5XTD and 9TI4, applying element-specific van der Waals exclusion with respect to protein and FMN atoms to obtain an accessible ensemble. A 16-nucleus Schulten–Wolynes calculation then provided singlet yields and their responses to magnetic field and dephasing.(17, 32) We combined these quantities in a composite score and examined its dependence on radical separation, structural input and kinetic parameters. We also tested how imposed changes in FMN–O2 separation affect yield variability. Excluding positions that overlap FMN removes the near-contact score maximum reported in an earlier version of this analysis; the accessibility-weighted score instead peaks several ångströms from the ring centre, at a separation that depends on the structural model.

## Results

### Structure-guided oxygen sampling and spin-response calculations

We generated 120,000 candidate O_2_ centroids around the FMN isoalloxazine ring in human complex I (PDB 5XTD) and removed positions that violated element-specific steric exclusion criteria with respect to both the protein and the FMN cofactor itself (**Fig. 1A,B**). Accessible configurations were grouped by their FMN–O_2_ centroid distance, r, to obtain a local acceptance fraction, A(r). This fraction measures geometric accessibility under the sampling procedure, not oxygen occupancy or binding probability (**Fig. 1E**). For each separation, a semiclassical Schulten–Wolynes calculation sampled the hyperfine fields of 16 FMN nuclei (four ¹□N and twelve ¹H) and propagated the two-electron radical-pair state under Zeeman, exchange and phenomenological dephasing terms (**Fig. 1C,D**). Reference rates were k_S_ = k_T_ = 1 × 10□s□¹ and k_STD_ = 3 × 10□s□¹. We calculated the absolute change in singlet yield between 0.05 and 50 mT and the change between low and high dephasing rates. Each response was normalised separately; their mean was multiplied by normalised accessibility to obtain the composite score H(r) (**Fig. 1F**). High-scoring sampled positions occupied a restricted part of the accessible FMN environment in the reference distance range (**Fig. 1G**). Exchange–hyperfine and dipolar–hyperfine comparisons were examined afterward as mechanistic diagnostics; they were not terms in H(r) (**Fig. 1H**).

**Figure 1.**
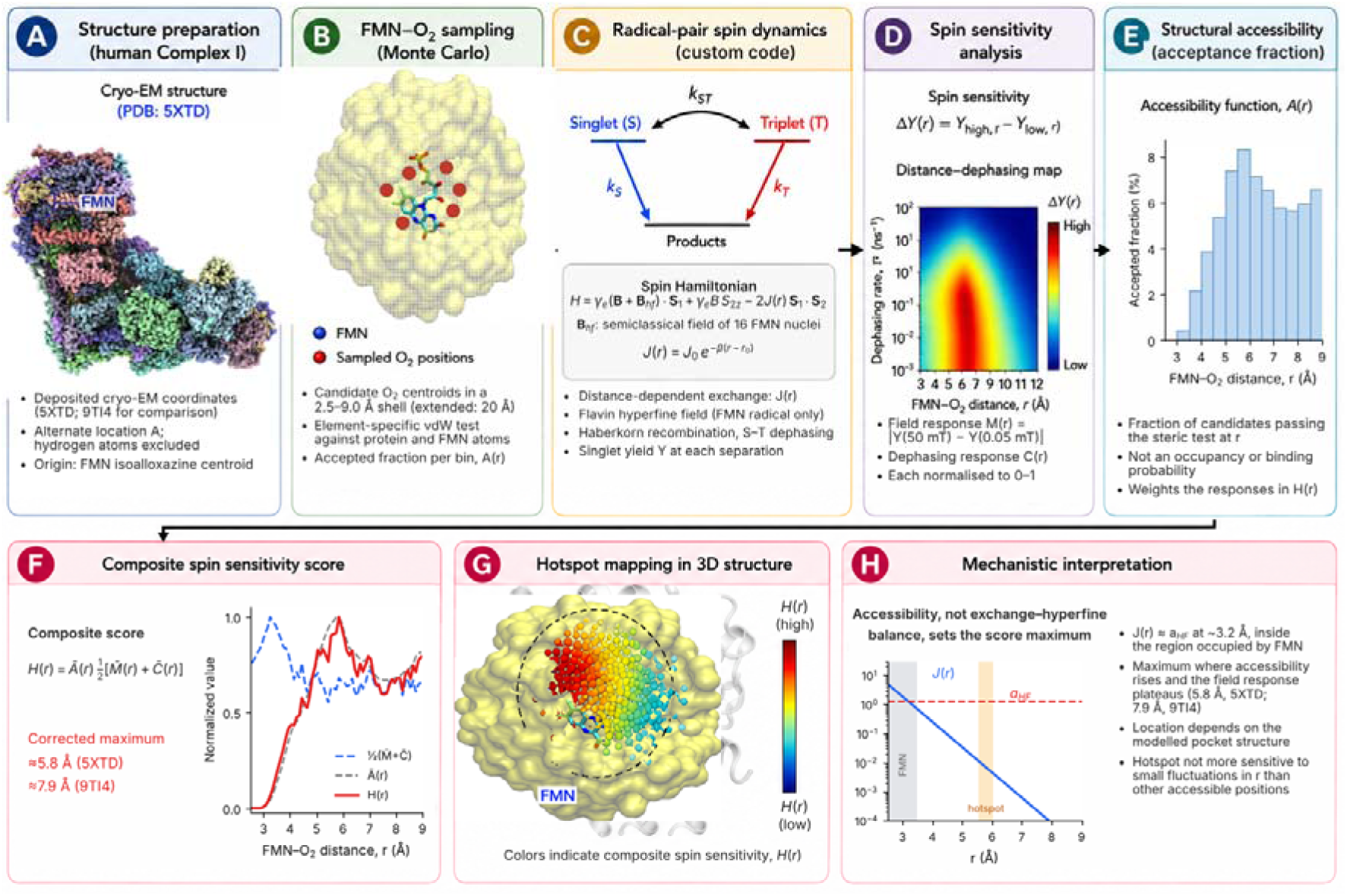
Computational workflow for the FMN–O_2_ radical-pair model. (A) Human complex I reference structure (PDB 5XTD) and the FMN-centred sampling region; deposited coordinates were used without modification. (B) Candidate oxygen centroids retained after steric filtering against protein and FMN atoms (schematic). (C) The two-electron spin Hamiltonian with a semiclassical hyperfine field from 16 FMN nuclei, distance-dependent exchange J(r), Haberkorn recombination and singlet–triplet dephasing. (D) Singlet reaction yield as a function of separation and dephasing (schematic); the field and dephasing responses are defined from singlet-yield differences. (E) The oxygen accessibility profile A(r), defined as the accepted fraction of trial centroids in each distance bin; it is not an equilibrium occupancy probability (5XTD, corrected steric test). (F) Composite score obtained from accessibility and the mean of separately normalised magnetic-field and dephasing responses (5XTD); the maximum lies near 5.8 Å. (G) High-scoring configurations mapped onto the structure (schematic). (H) Exchange coupling and the effective FMN hyperfine field: J(r) equals a_HF_ near 3.2 Å, inside the region occupied by FMN (grey), whereas the score maximum (orange) lies where accessibility rises; neither exchange–hyperfine balance nor singlet–triplet mixing is a component of the score.

### Distance dependence of the reference spin calculation

Exchange coupling in the model decreased exponentially with increasing FMN–O_2_ centroid separation, while the reference root-sum-square flavin hyperfine field was 1.23 mT (anionic FMN•□ parameter set; see Methods) (**Fig. 2A**). With FMN atoms included in the steric test, no accessible configurations occurred below 3.0 Å and only 0.2% of candidates between 3.0 and 3.3 Å were accepted; accessibility reached its maximum near 5.8 Å. The dephasing-response metric was greatest near contact and declined with distance (**Fig. 2B**). By comparison, the time-domain singlet–triplet mixing-amplitude proxy changed little across the interval (0.216 ± 0.00003); its dominant hyperfine/Zeeman contribution did not depend on r. This near-constant mixing metric should not be read as a distance-independent singlet–triplet conversion rate. The calculated singlet yield varied with both separation and the imposed dephasing rate (**Fig. 2C**). For the reference composite score, the largest value occurred at 5.85 Å, and the middle 95% of top-decile configurations spanned 5.56–6.04 Å (**Fig. 2D**). This interval describes the selected configuration distribution, not a confidence interval for an experimentally measured optimum.

**Figure 2.**
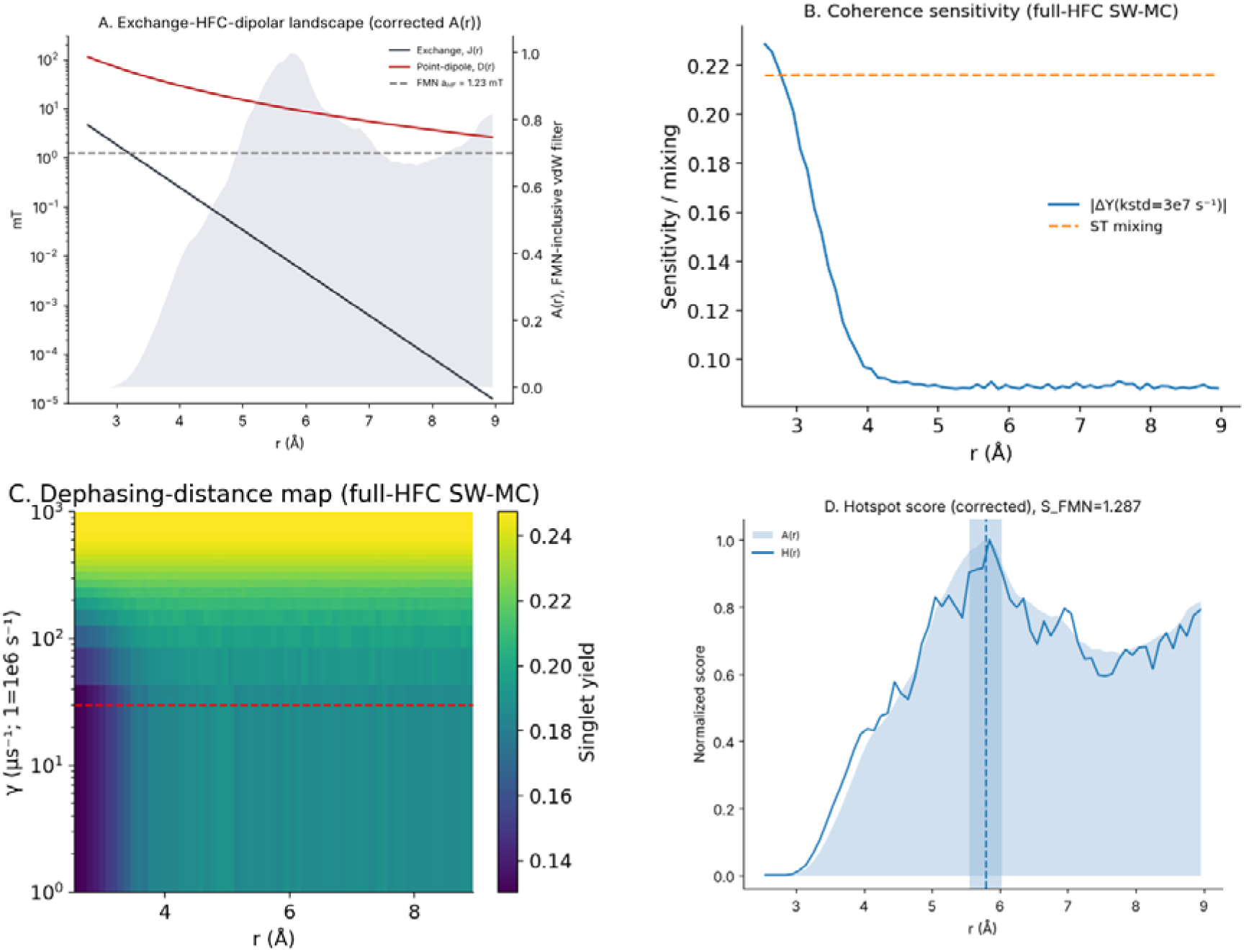
Distance-dependent quantities in the FMN–O_2_ radical-pair model. (A) Exchange J(r), collinear point-dipole coupling D(r), reference FMN hyperfine field (1.23 mT) and the steric accessibility profile from 5XTD. Exchange parameters are J0 = 5.0 mT, r0 = 2.5 Å and β = 2.0 Å□¹. (B) Dephasing-response sensitivity and the time-domain singlet– triplet mixing-amplitude proxy. The mixing measure is an amplitude, not a transition rate. (C) Calculated singlet yield versus separation and dephasing rate. (D) Composite score H(r); shading marks the central 95% of distances among the highest-scoring 10% of accepted configurations, and the dashed line indicates their mean. Exchange–hyperfine balance and the mixing proxy were calculated separately. Accessibility excludes positions overlapping FMN. Panels A and D use the corrected accessibility; panels B and C are unchanged from the previous version.

### Contributions to the composite-score maximum

Figure 3 separates the inputs to the composite score. The exponential exchange term approaches the 1.23 mT effective hyperfine scale over a short distance range (**Fig. 3A**). Accessibility is negligible below about 3.5 Å and rises to a maximum near 5.8 Å, whereas the dephasing-response component is largest near contact and falls with separation (**Fig. 3B**). Because the dephasing response is greatest where FMN leaves no room for oxygen, it contributes little to the accessibility-weighted score. Its combination with the normalised magnetic-field response, which is suppressed near contact and approaches a plateau beyond about 5 Å, yields a score maximum at 5.85 Å in the reference calculation (**Fig. 3C**). Selecting the highest-scoring 10% of accessible configurations gives a mean distance of 5.80 Å and a standard deviation of 0.14 Å, with 2.5th–97.5th percentiles of 5.56–6.04 Å (**Fig. 3D**). The grid maximum was the same (5.85 Å) for five independent sampling seeds, and the top-decile mean had a standard deviation of 0.03 Å across seeds; because the grid position varied between 5.5 and 6.0 Å in the alternative spin models (Methods), we report the top-decile distribution as the primary measure. The approximately constant singlet–triplet mixing-amplitude proxy provides no separate evidence for a mixing optimum at these separations.

**Figure 3.**
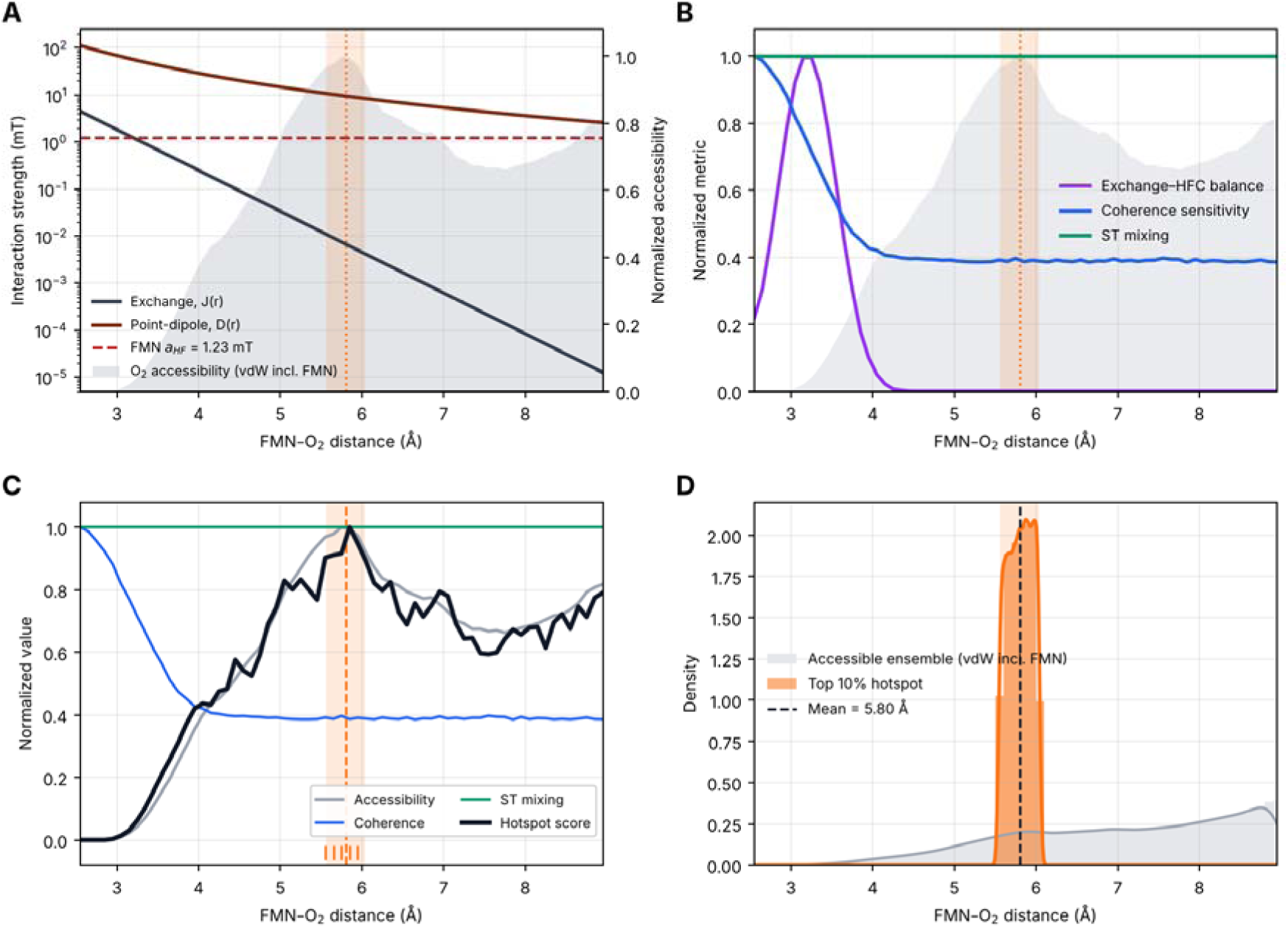
Contributions to the composite-score maximum. (A) Model exchange and collinear point-dipole couplings relative to the 1.23 mT effective FMN hyperfine field. (B) Steric accessibility, dephasing response and spin diagnostics as functions of separation. (C) Composite score from accessibility and normalised field and dephasing responses; tick marks show the highest-scoring sampled configurations. (D) Distribution of the top-decile accepted configurations. In the 2.5–9.0 Å calculation, the mean is 5.80 Å, the standard deviation is 0.14 Å and the 2.5th–97.5th percentile interval is 5.56–6.04 Å. Spin-response profiles are those of the previous version; accessibility is corrected. The composite-score maximum is not a maximum in raw magnetic-field sensitivity.

### Dependence of the score maximum on structure, model parameters and distance domain

The location of the score maximum depended on the structural model. With the independently determined 9TI4 structure (2.66 Å resolution), the top-decile mean was 7.86 Å (SD 0.07 Å; 2.5th–97.5th percentiles, 7.74–7.98 Å) (Table 1; **Supplementary Figs S3 and S5**). In 9TI4, accessibility rose more slowly with separation and reached its maximum near 7.8 Å; Asn116 and Asn244 of the NDUFV1 subunit lie 3.3 and 3.5 Å from the isoalloxazine centre, compared with 4.5 and 7.4 Å in 5XTD. Within each structure, the location was insensitive to the model choices tested. Across three dephasing models, two flavin hyperfine parameter sets and four definitions of the response metrics, the top-decile mean in a re-implementation of the spin model ranged from 5.4 to 5.8 Å in 5XTD and from 7.9 to 8.2 Å in 9TI4. Varying the O_2_ probe radius (1.40–1.60 Å), scaling the FMN van der Waals radii by 0.8–1.0, including hydrogen atoms or using Rowland–Taylor radii changed these means by at most 0.45 Å (**Supplementary Fig. S5**).

**Table 1.** Composite-score peak and distance distribution of top-decile configurations after exclusion of FMN-overlapping positions. Rows for 2.5–9.0 Å pool 2,000,000 candidate O2 positions and the original spin-response profiles; extended-range rows use 1,300,000 candidates and the re-implemented spin model (Methods). The final column gives the 2.5th–97.5th percentiles of selected distances, not confidence intervals for the peak. In the exchange-only extended row, the highest grid value lies at the domain boundary (*) and does not define a localized maximum. The near–contact peaks (2.75–3.35 Å) listed in the previous version were obtained with a steric test that omitted FMN atoms and are superseded.

| **Domain** | **Model** | **Peak (Å)** | **Mean (Å)** | **SD (Å)** | 95% range (Å) |
|---|---|---|---|---|---|
| 2.5–9.0 Å, 5XTD (main text) | exchange-only | 5.85 | 5.80 | 0.14 | 5.56–6.04 |
| 2.5–9.0 Å, 9TI4 | exchange-only | 7.85 | 7.86 | 0.07 | 7.74–7.98 |
| 2.5–20 Å, 5XTD | exchange-only | 19.35* | 19.62 | 0.24 | 19.27–19.99 |
| 2.5–20 Å, 5XTD | exchange + dipolar | 6.05 | 8.18 | 1.43 | 5.39–10.13 |
| 2.5–20 Å, 9TI4 | exchange + dipolar | 7.85 | 9.53 | 1.82 | 6.81–13.38 |

We then extended the analysis to 2.5–20 Å with 1,300,000 candidate positions, using either exchange coupling alone or exchange plus a collinear point-dipole term (Table 1; Supplementary Notes 1 and 2). With exchange alone, the score formed a broad plateau beyond about 5 Å, and its highest grid value lay at the 20 Å boundary in both structures, so no localized maximum could be defined. With the dipolar term, the maximum lay at 6.05 Å in 5XTD and 7.85 Å in 9TI4, and the top-decile distributions were broad (means 8.18 and 9.53 Å). No near-contact maximum appeared in any extended-range calculation.

### Effect of imposed separation fluctuations on singlet yield

We next varied FMN–O_2_ centroid separation around sampled baseline positions and interpolated the reference singlet-yield curve (**Fig. 4A**). Gaussian perturbations increased the dispersion of calculated singlet yields, with smaller changes in their mean (**Fig. 4B**). Because the corrected high-scoring interval (5.50–6.00 Å in 5XTD) lies on the plateau of the yield curve and the long-distance comparison region used previously (≥6.5 Å) would overlap it, configurations in the top-decile interval were compared with all other accessible configurations. In 5XTD, within-configuration yield standard deviations in the high-scoring interval were comparable to those elsewhere at σ = 0.10 and 0.25 Å (ratios 1.23 and 0.77; both groups near the noise level of the yield curve), smaller at σ = 0.50 and 0.75 Å (ratios 0.39 and 0.74), and larger only at σ = 1.00 Å (ratio 1.60), when perturbations reached the steep part of the yield curve below 4.5 Å (**Fig. 4C**). In 9TI4, the ratio fell from 1.44 at σ = 0.10 Å to 0.23 at σ = 1.00 Å. Yield variance divided by the realised variance of perturbed separation was largest at the sparsely accessible inner edge of the pocket (3.45 Å in 5XTD and 4.35 Å in 9TI4) rather than within the high-scoring interval (**Fig. 4D**). The calculation describes sensitivity to a prescribed geometric perturbation; no protein-dynamics trajectory or experimentally measured fluctuation amplitude was used.

**Figure 4.**
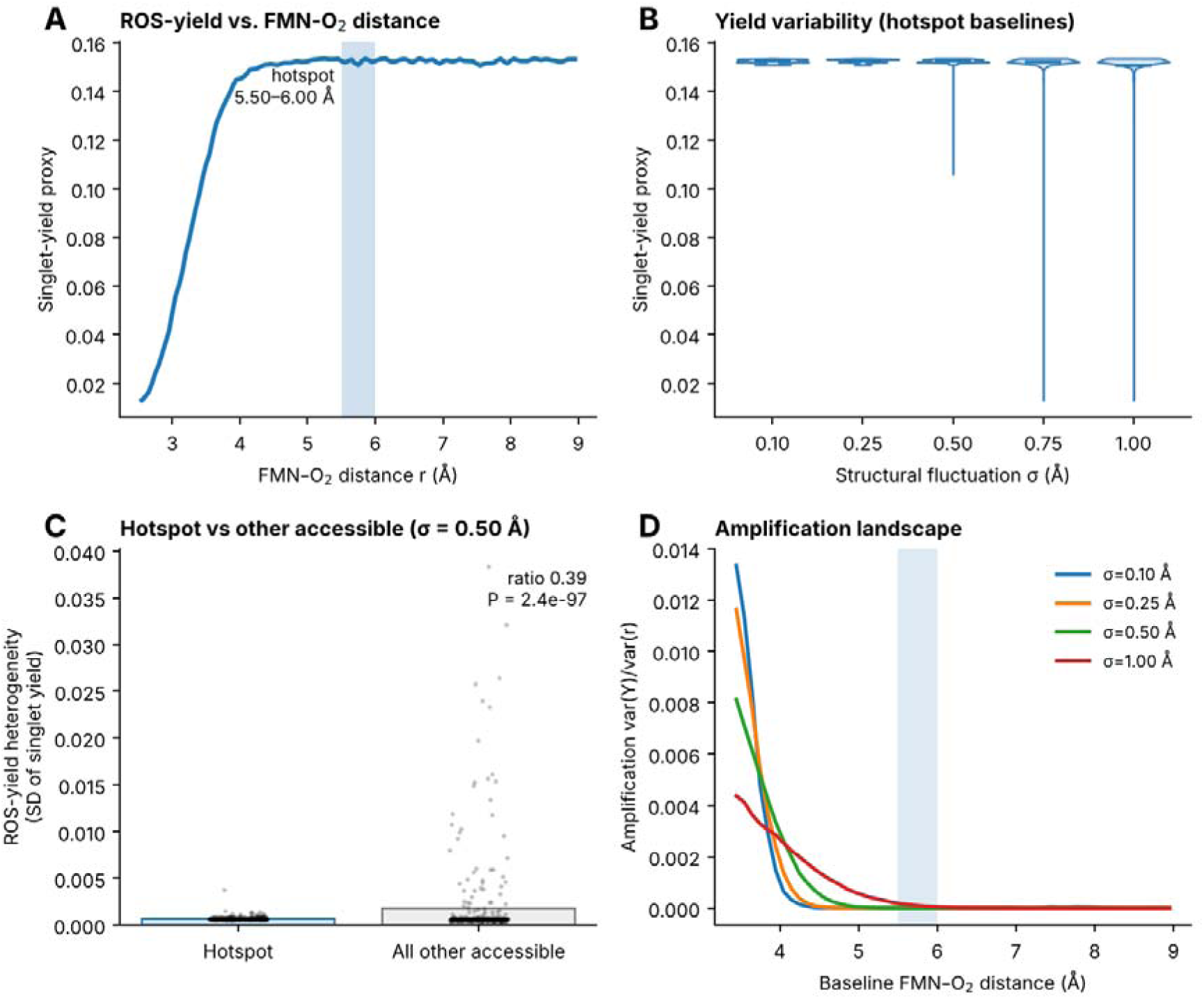
Response of modelled singlet yield to imposed FMN–O_2_ separation fluctuations. (A) Singlet yield against centroid separation, with the reference top-decile composite-score region shaded. (B) Simulated yield distributions after Gaussian separation perturbations with σ = 0.10, 0.25, 0.50, 0.75 and 1.00 Å and boundary clipping at 2.5 and 9.0 Å. (C) Within-configuration yield standard deviations in the top-decile interval and in all other accessible configurations. Error bars indicate approximate 95% confidence intervals for group means; group comparisons used two-sided Mann–Whitney U tests. (D) Ratio Var(Y)/Var(r) plotted against baseline separation. This response measure describes imposed geometric perturbations, not observed protein motion. The singlet-yield curve is that of the previous version; baselines are drawn from the corrected accessible ensemble of 5XTD, and panel B shows baselines in the top-decile interval.

### Dependence on recombination and spin-dephasing rates

The reference rates were k_S_ = k_T_ = 10□s□¹ and k_STD_ = 3 × 10□s□¹; they were not measured for the FMN–superoxide pair in complex I. We carried out three sensitivity tests (**Fig. 5**). In the first, all three rates were set equal and swept together from 10 to 10□s□¹. In the second, k_STD_ alone was varied while k_S_ and k_T_ stayed at 10□s□¹. In the third, k_S_ = k_T_ was varied between 10 and 10□s□¹ with k_STD_ held at 3 × 10□s□¹. The joint sweep is a separate parameter family and must not be confused with the reference calculation.

**Figure 5.**
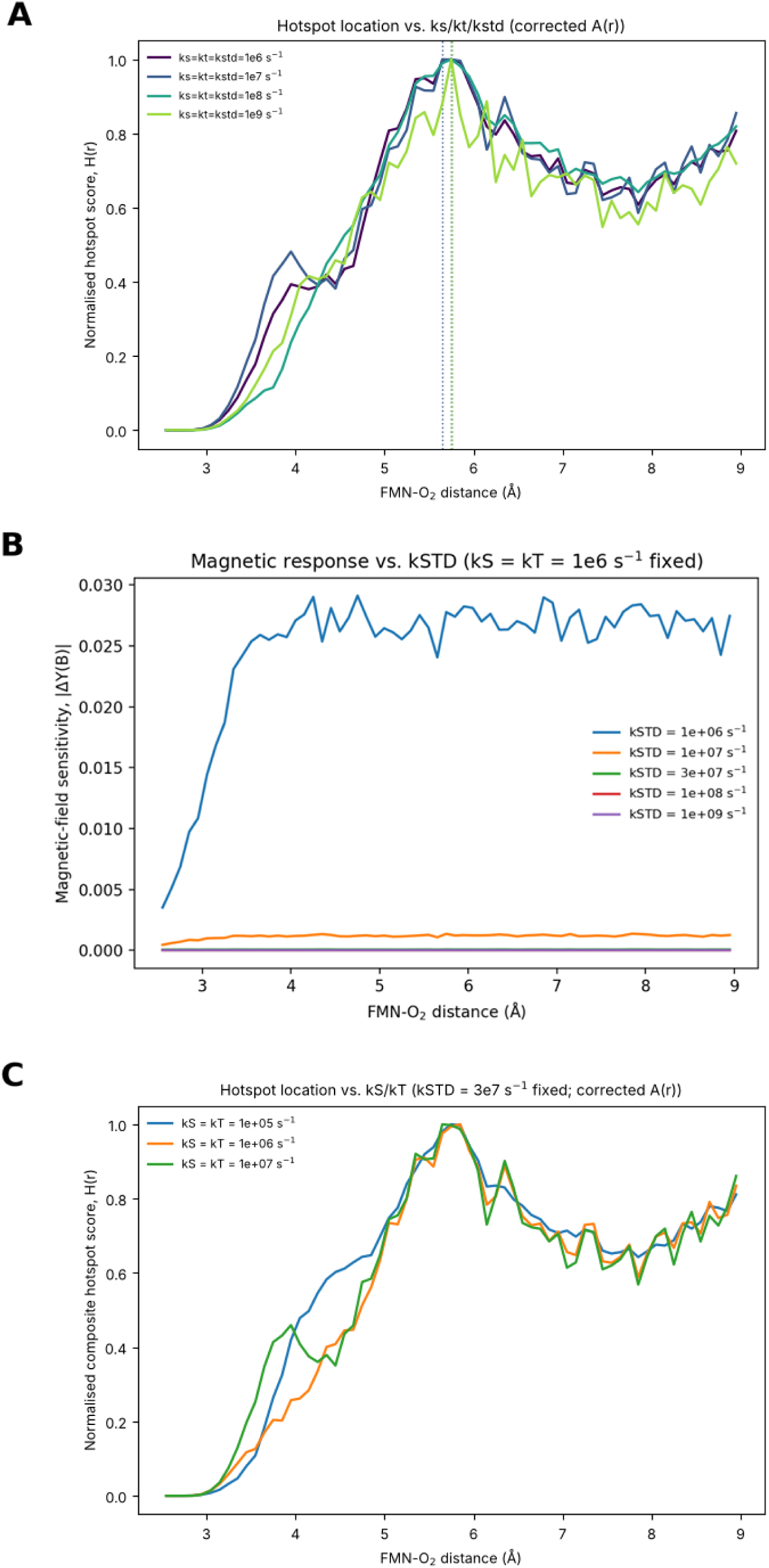
Dependence on recombination and dephasing rates. (A) Composite score with k_S_ = k_T_ = k_STD_ jointly set to 10□, 10□, 10□ or 10□s□¹. These are not the reference rate values. (B) Raw |ΔY(B)| for varying k_STD_ at fixed k_S_ = k_T_ = 10□s□¹. (C) Composite scores for k_S_ = k_T_ of 10□, 10□and 10□s□¹ with k_STD_ fixed at 3 × 10□s□¹. Panels A and C were recalculated with the corrected accessibility using the re-implemented spin model (Methods); panel B is unchanged. A stable normalised score peak does not imply a large absolute field response.

Across the joint sweep, composite-score peaks lay between 5.65 and 5.75 Å and top-decile means between 5.67 and 5.72 Å in 5XTD (7.75–8.15 Å and 7.81–8.05 Å in 9TI4; re-implemented spin model, Methods) (**Fig. 5A**). Changing k_STD_ alone strongly suppressed the absolute field response: its maximum fell to 4.7% of the 10□s□¹ result at 10□s□¹ and to 0.26% at the reference 3 × 10□s□¹ (**Fig. 5B**). When kS = kT alone was varied, the score peak remained at 5.65–5.85 Å and the mean at 5.73–5.74 Å in 5XTD (7.75 Å and 7.83–7.98 Å in 9TI4) (**Fig. 5C**). The score’s location was less sensitive to these rates than the magnitude of the underlying magnetic-field effect. In particular, a well-defined peak in normalised H(r) remains possible even when absolute |ΔY(B)| is almost extinguished.

The raw magnetic-field response, |ΔY(B)|, did not show a narrow optimum. In the original 2.5–9.0 Å sweep, the largest sampled values occurred at 4.75 Å for k_STD_ = 10□s□¹ and 7.85 Å for the higher rates (**Supplementary Fig. S2**). A subsequent calculation with more spin samples and the range extended to 20 Å showed why those apparent maxima are misleading (**Supplementary Note 2, Fig. S4**): |ΔY(B)| was suppressed near contact and approached a broad, nearly flat response at longer separation. The exchange-plus-dipolar calculation also showed a weak, broad excess at intermediate long distances under low dephasing, but no narrow second peak. The 4.75 and 7.85 Å positions therefore should not be treated as distinct magnetic-response optima. With FMN-overlapping positions excluded, the rise in accessibility and the onset of the field-response plateau chiefly determine the location of the composite-score maximum.

Simulations of other flavoproteins found O_2_ unbinding times of 12–43 ns in electron-transfer flavoprotein and localized O_2_ and O_2_^•□^ in cryptochrome.(42, 43) The present accessibility profile does not calculate residence time. Cryptochrome-based calculations estimated a rotational correlation time of 0.99 ± 0.33 ps and a spin-rotational relaxation time of about 0.8 ns for free superoxide, with slower rotation in some protein-associated configurations.(41) These observations motivated the rate sweep in Fig. 5. Here k_STD_ is an imposed radical-pair dephasing rate, not a measured value for complex I.(40)

## Discussion

The corrected composite-score calculation places its highest accessibility-weighted spin response several ångströms from the isoalloxazine centre, not at near contact. Near contact the dephasing response is largest, but FMN itself occupies almost all of that space; positions there were counted as accessible in the previous version of this analysis because the steric test omitted FMN atoms. With these positions excluded, the score maximum lies where accessibility rises and the raw magnetic-field response approaches its plateau. Examining these quantities separately distinguishes the geometries favoured by the structural model from those showing the largest absolute field response.

The location of that maximum depends on the structural input. The top-decile mean distances were 5.80 Å for 5XTD and 7.86 Å for 9TI4 (Table 1; **Supplementary Figs S3 and S5**).(22) The agreement between the two structures reported previously (3.28 and 3.25 Å) reflected the shared omission of FMN atoms, which dominated both accessibility profiles; once it is corrected, differences in the modelled side chains around the flavin determine where the score is highest. Because the result was stable to the spin-model and steric-parameter choices within each structure, uncertainty in the pocket geometry, rather than in the spin model, limits how precisely a preferred oxygen-approach distance can be stated. Over 2.5–20 Å, the exchange-only score had no localized maximum.

The perturbation calculations do not identify the high-scoring configurations as especially sensitive to small changes in FMN–O_2_ separation. The largest yield response to imposed fluctuations occurred at the sparsely accessible inner edge of the pocket, where the singlet yield changes steeply with distance (**Fig. 4**). In the proposed FMN–superoxide reaction scheme, the singlet-selective channel is associated with H_2_O_2_ formation.(33) The calculated dependence on geometry suggests an experiment in which peroxide/superoxide product branching is measured while the FMN-pocket environment is changed.

The flavin-localization studies provide a useful physical setting for these predictions. Molecular-dynamics simulations of electron-transfer flavoprotein and cryptochrome have found oxygen near flavin groups and longer-lived superoxide configurations in some protein pockets.(42, 43) Applying similar simulations to complex I would show which high-scoring approach geometries are visited during oxygen motion and how the resulting superoxide radical moves after electron transfer. Those trajectories could be paired with spin calculations for the same configurations.

The rate sweeps also identify conditions in which the modelled response is strongest. Varying the dephasing rate changed the magnetic-field response far more than the position of the score maximum (**Fig. 5**). Protein-associated superoxide can rotate more slowly than the free radical in model systems.(40, 41) Measuring superoxide motion in the FMN pocket would permit kinetic parameters specific to complex I.

A next step is to combine oxygen and superoxide trajectories with orientation-dependent exchange and dipolar coupling and measurements of field-dependent product branching. The different results obtained from two complex I structures indicate that such tests should use structural ensembles, for example from molecular-dynamics simulations, rather than a single model. This approach, and future work, would assess whether the configurations selected by the structural map contribute to a detectable change in peroxide or superoxide yield.

The corrected calculations show that the accessibility-weighted spin-response score of the complex I flavin site has no near-contact maximum and that its location is governed mainly by pocket geometry, differing between two cryo-EM structures. They correct the near-contact feature reported previously and provide distances and observables for experiments on radical-pair reactivity at the complex I flavin site.

## Methods

### Structural model and FMN-centred oxygen sampling

The reference structure was the cryo-electron microscopy model of human mitochondrial complex I (PDB 5XTD). FMN and neighbouring protein coordinates were taken from the deposited structure. The centre of the FMN isoalloxazine ring defined the origin of the geometric FMN–O_2_ distance coordinate.(23, 24)

We generated 120,000 candidate O_2_ centroid positions within the modelled 2.5–9.0 Å domain around FMN.(25) A candidate was accepted only when its distance from the nearest heavy atom of each element (C, N, O, S, P or Fe), including the atoms of the FMN cofactor itself, exceeded the sum of that element’s van der Waals radius and an O_2_ probe radius of 1.52 Å.(26) This element-specific hard-sphere test rejects more positions than the single-distance exclusion used in an earlier version of the calculation. In the previous version of this analysis, FMN atoms were omitted from the test; 99–100% of the positions it accepted within 3.3 Å of the ring centre overlapped FMN atoms (median distance to the nearest FMN atom, 1.2–1.35 Å). All results reported here use the corrected test. Fe, which has no Bondi radius, was assigned 2.00 Å. Accepted centroids formed the geometric accessibility ensemble. The separation r was measured between each O_2_ centroid and the FMN isoalloxazine centre, and was used as the input to the exchange model. It is a structural proxy, not a calculated electron–electron distance. The acceptance procedure does not give an equilibrium oxygen occupancy, diffusion pathway, binding free energy or residence time. It tests positions against a fixed structure, without simulating O_2_ trajectories or the motion of newly formed superoxide.(27, 28, 42, 43)

### Structure-derived oxygen-accessibility profile

For each distance bin, A(r) was the number of accepted O_2_ centroids divided by the total number of candidates sampled in that bin. This acceptance fraction avoids the radial shell-volume weighting that would arise from using raw accepted counts. A bin containing few candidates nevertheless has greater sampling uncertainty; the extended-domain calculations were repeated with a larger candidate ensemble to assess this effect (Supplementary Notes 1 and 2). The binned profile was smoothed, interpolated onto the spin-calculation grid and normalised to its maximum.(29, 30)

The accessibility profile was normalised to its maximum value:

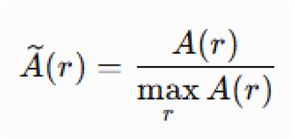

Accordingly, represents the relative geometric accessibility of candidate O_2_ configurations at a given FMN–O_2_ separation.(*31*) The accessibility profile was derived independently of the radical-pair spin model and was incorporated only during the subsequent composite spin-sensitivity analysis.

### FMN–O2 radical-pair model

We used a two-electron model with an FMN semiquinone radical and superoxide.(32) The initial spin state was the maximally mixed triplet-subspace density operator

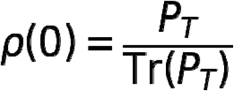

where PT denotes the triplet-subspace projector.(15) We adopted this initial state as a modelling assumption based on spin conservation during electron transfer from reduced FMN to triplet-ground-state O_2_.(33)

Triplet-ground-state oxygen motivates a triplet-correlated initial pair in the proposed reaction scheme.(33) The spin state after electron transfer has not been measured in the complex I FMN pocket, so we did not treat the initial triplet population as experimentally established.

In the proposed product-branching scheme, a triplet-channel reaction releases superoxide (O ^•□^), whereas a singlet-associated second electron transfer can lead to hydrogen peroxide (H_2_O_2_).(33) We therefore use the modelled singlet-reaction yield as a proxy for the H_2_O_2_-associated channel under this scheme. Its complement describes the other spin-selective channel only within the two-channel model; it is not an independently measured superoxide yield. Results may change for other initial spin-state populations. The time-dependent singlet population was calculated as

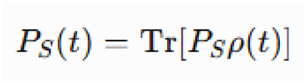

where is the time-evolved radical-pair density matrix. (*32*) The radical-pair spin Hamiltonian included electron Zeeman, effective flavin hyperfine, and distance-dependent exchange interactions:

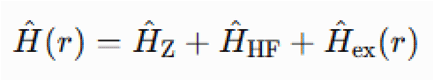

The effective hyperfine interaction associated with the FMN radical was represented by an effective coupling strength of

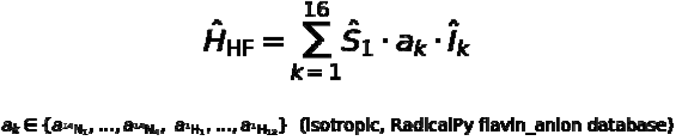

The flavin hyperfine field was calculated with the Schulten–Wolynes (SW) semiclassical approximation using the isotropic coupling constants of four ¹□N and twelve ¹H nuclei from the RadicalPy molecule database. Nuclear spins were represented by independently oriented classical vectors of length √[I(I+1)] and resampled for each Monte Carlo draw. The hyperfine-weighted sum of these vectors acted on the flavin electron spin. There were 400 draws per distance bin in the main calculation and 80 per bin in the illustrative dephasing-distance map. Since the nuclear spins are sampled classically, the propagated Liouville space contains only the two electron spins. The resulting effective root-sum-square hyperfine field was 1.23 mT. These constants correspond to the anionic flavin radical (FMN•) entry of the database; repeating the composite-score calculation with the neutral semiquinone (FMNH•; four ¹□N and seventeen ¹H nuclei, 1.108 mT) changed the top-decile mean by less than 0.05 Å. Haberkorn recombination rates were set to k_S_ = k_T_ = 10□s□¹ and the reference singlet– triplet dephasing rate to k_STD_ = 3 × 10□s□¹. These are model inputs, not complex I measurements.

### Distance-dependent exchange interaction

Exchange coupling between the FMN semiquinone and superoxide radical electrons was represented using an exponential distance-decay model:

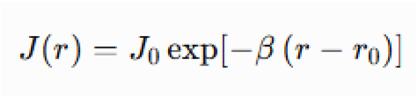

where *r* is the geometric FMN–O_2_ separation coordinate, *J*_0_ is the exchange coupling assigned at the reference distance *r*_0_, and β is the exponential distance-decay constant. (*34*)

The model parameters were

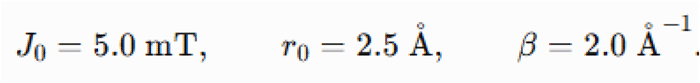

We used J0 = 5.0 mT, r0 = 2.5 Å and β = 2.0 Å ¹ in the exponential exchange model. The chosen values provide a distance-dependent test case; they have not been fitted to exchange-coupling measurements for the FMN–superoxide pair.(35)

The exchange interaction was incorporated into the spin Hamiltonian as

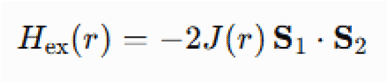

where S_1_ and S_2_ are the electron-spin operators associated with the FMN semiquinone and superoxide radicals, respectively. These parameters define a phenomenological distance-dependent exchange model and should not be interpreted as experimentally measured exchange couplings for the FMN–superoxide radical pair. The model was used to represent the expected rapid attenuation of exchange interactions with increasing radical separation and to examine the resulting spin-dynamic regimes across the structure-derived FMN–O_2_ landscape. (*36*)

### Radical-pair spin dynamics and singlet-reaction yield

A custom Python/NumPy implementation propagated the electron-spin density matrix using Schulten–Wolynes Monte Carlo hyperfine sampling. RadicalPy supplied the molecular hyperfine constants but was not used as the spin-propagation backend.(17, 37)

Environmental spin dephasing was represented phenomenologically through a dephasing rate, γ. The evolution of the radical-pair density matrix can be expressed generally as

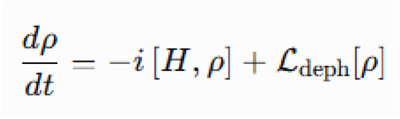

where *H* is the radical-pair spin Hamiltonian and *L*_dp_ represents the spin-dephasing contribution.(*38*) For each distance and spin-environment condition, the singlet-selective reaction yield was calculated from the time-dependent singlet population:

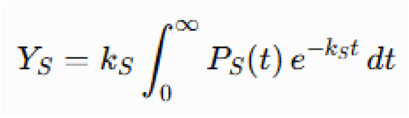

Here kS is the singlet recombination rate. The reference parameters were kS = kT = 10□s□¹ and k_STD_ = 3 × 10□s□¹.(39) This model quantity is a singlet-selective branch yield under the stated spin Hamiltonian and kinetic assumptions; it is not the total cellular ROS yield.

The relaxation parameter k_STD_ was chosen for sensitivity analysis; it was not derived from oxygen rotational motion. Specifically, the spin calculation contains no explicit rotational correlation time for superoxide (O_2_ ^•□^) and no separate spin-rotational relaxation time. The reciprocal 1/k_STD_ is therefore only the timescale of the imposed dephasing term. Rotational and spin-relaxation times reported for superoxide in cryptochrome models should not be assigned to complex I without a separate dynamical calculation.(40, 41)

### Dephasing–distance singlet-yield landscape

To characterise the dependence of radical-pair reactivity on molecular separation and environmental spin dephasing, singlet reaction yields were evaluated across a two-dimensional grid of FMN–O_2_ separation distances and dephasing rates.

For each combination of r and (γ), the singlet reaction yield was calculated as

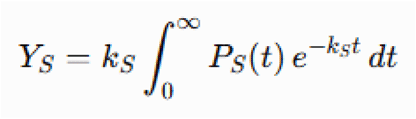

Dephasing rates were sampled on a logarithmic grid. Differences in singlet yield across that grid describe sensitivity to the imposed dephasing parameter; they do not by themselves quantify a measured coherence lifetime.(39)

### Magnetic-field and dephasing sensitivities

Magnetic-field sensitivity was the absolute difference in the computed singlet yield at 50 mT and approximately 0.05 mT. At each separation and rate set, both field calculations reused the same Monte Carlo nuclear-spin orientations (common random numbers, obtained with an identical per-bin seed). This reduced stochastic error in their difference. The field comparison is a model perturbation spanning three orders of magnitude, not a direct estimate of sensitivity to a small change in geomagnetic field strength.

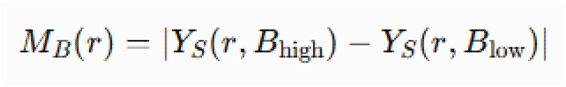

Dephasing sensitivity was quantified as the absolute difference in singlet reaction yield between the low- and high-dephasing conditions:

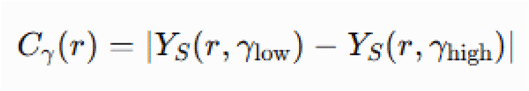

The magnetic-field and dephasing sensitivity profiles were independently normalised to the range 0–1:

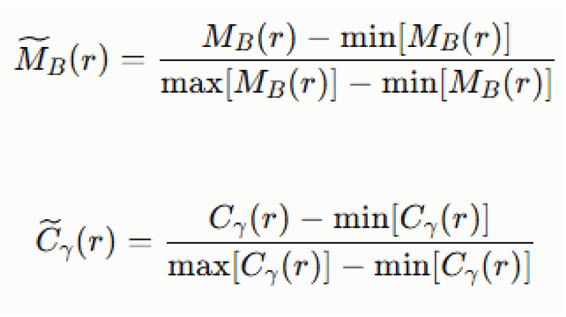

Each response function measures a change in calculated singlet yield under a specified model perturbation. The dephasing response is not a direct measure of quantum coherence.(15)

### Composite spin-sensitivity score

A composite spin-sensitivity score was calculated to identify FMN–O_2_ configurations that were both structurally accessible and dynamically responsive to changes in the spin environment.

The composite score was defined as

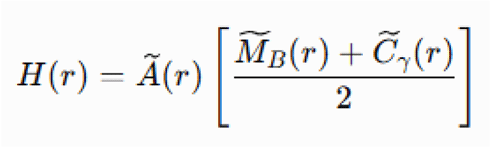

where Ã(r)is the normalised structure-derived oxygen accessibility, M̃ × B(r) is the normalised magnetic-field sensitivity, and C̃ × γ r is the normalised dephasing sensitivity. (*29, 30*)

The two normalised response terms were given equal weight as a modelling choice, and their mean was multiplied by normalised accessibility. The score was then divided by its maximum to give H_norm_(r). These operations rank the sampled geometries; they do not preserve the absolute magnitude of magnetic-field sensitivity.

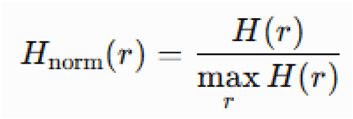

Exchange–hyperfine balance and singlet–triplet mixing were evaluated separately as mechanistic diagnostic quantities and were not included in the definition or calculation of the composite spin-sensitivity score.

### Identification of spin-sensitive hotspot configurations

Each sterically accessible FMN–O_2_ configuration was assigned the normalised composite score corresponding to its separation coordinate. Accessible configurations were ranked according to H_norm_(r), and the highest-scoring 10% of the accessible ensemble were operationally classified as spin-sensitive hotspot configurations.

The threshold was the 90th percentile of the score among accepted configurations. We report the selected distances by their mean, standard deviation, and 2.5th–97.5th percentiles. The percentile interval is a descriptive range of configurations rather than a confidence interval for a fitted parameter.

The 10% threshold is a ranking convention. Its location and breadth can change when the distance domain or the response weighting changes.

Hotspot enrichment was quantified using

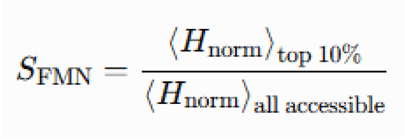

where the numerator is the mean normalised composite score of the highest-scoring 10% of accessible configurations and the denominator is the corresponding mean across the complete structurally accessible ensemble.

### Kinetic-parameter sensitivity sweep

For the joint kinetic sweep, k_S_ = k_T_ = k_STD_ was set successively to 10□, 10□, 10□and 10□s□¹, and the composite-score calculation was repeated at each setting. This sweep does not contain the reference combination k_S_ = k_T_ = 10□s□¹, k_STD_ = 3 × 10□s□¹. For each case, we recorded the highest-scoring grid point and the mean, standard deviation and percentile bounds of the top-decile distance distribution. Separate parameter sweeps are described below.

### Structural ensemble reconstruction and additional single-parameter/dipolar sensitivity analyses

The oxygen ensemble was regenerated from PDB 5XTD with the same element-specific exclusion rule and O_2_ probe radius (1.52 Å). In the reference distance domain, the alternative 9TI4 structure was also analysed without changing the physical parameters. 9TI4 coordinates were read from the mmCIF file, retaining alternate location A where present; hydrogen atoms were excluded from the reference steric test. The single-parameter sweeps and 2.5–20 Å dipolar comparison used the reconstructed 5XTD ensemble.

For the k_STD_-only sweep, k_S_ = k_T_ remained at 10□s□¹ while k_STD_ ranged from 10 to 10□s□¹. Absolute field sensitivity was calculated as the singlet-yield difference between 50 and approximately 0.05 mT. For the k_S_ = k_T_-only sweep, k_STD_ remained at 3 × 10□s□¹ and recombination rates were set to 10□, 10□ and 10□s□¹. We recalculated the composite score for each parameter set.

For the extended-distance calculation, 1,300,000 candidates were sampled over 2.5–20 Å (a 120,000-candidate run was used in the previous version) (Supplementary Notes 1 and 2). A point-dipole coefficient, D(r) = μ_0_g²μ_B_²/(4πhr³), was used in an axial secular term D(r)[3S_1z_S_2z_ − S_1_·S_2_]. The earlier implementation used the opposite overall sign of this term; the corrected operator is D(r)[S□S□− 3S□zS□z], and with the corrected steric test the sign changed the top-decile statistics by at most 0.05 Å. The calculation assumed the FMN–O_2_ vector to be collinear with the field axis. This is a fixed-orientation approximation, not a tensor orientation average. The exchange-only and exchange-plus-dipolar scores used k_S_ = k_T_ = 10□s□¹ and k_STD_ = 3 × 10□s□¹.

### Independent verification and sensitivity of the corrected accessibility calculation

Because the original spin-propagation code was not available for this revision, the corrected reference results combine the distance-dependent spin-response profiles of the original calculation with the recalculated accessibility. The dephasing response, the singlet-yield curve and the composite score of the previous version were digitized from its reference-calculation figures, and the normalised magnetic-field response was recovered as M (r) = s·H(r)/Ã(r) − C(r), with s fixed by the 0–1 normalisation (the recovered profile spanned 0.004–1.000). Recombining these profiles with the uncorrected accessibility reproduced the previous results for 5XTD (grid peak 3.25 Å; top-decile mean 3.24 Å, SD 0.24 Å, 2.5th– 97.5th percentiles 2.81–3.61 Å; previously 3.25 Å, 3.28 Å, 0.24 Å and 2.77–3.64 Å) and the previous perturbation contrast at σ = 0.50 Å (72-fold; previously 66-fold). The extended-domain, rate-sweep and spin-model sensitivity calculations used a re-implementation of the spin model, described below.

To check the steric calculation independently of random sampling and tree-based neighbour searches, accessibility was recomputed on a regular 0.1 Å cubic grid (2,988,301 points in the 2.5–9.0 Å shell) by brute-force evaluation of all atom–point distances, using a separately written coordinate parser. The grid and Monte Carlo calculations agreed (accessible fractions 5.98% and 5.96% for 5XTD) and gave the same top-decile distributions. Sensitivity to the steric parameters was assessed with 2,000,000 candidates per structure (five independent seeds), varying the O_2_ probe radius (1.40, 1.52 and 1.60 Å), scaling the FMN van der Waals radii by 0.8 or 0.9, including hydrogen, magnesium and zinc atoms, and replacing the Bondi radii of C, N, O and S with Rowland–Taylor values. Spin-model sensitivity used singlet– triplet dephasing, isotropic random-field relaxation of both electron spins and longitudinal (z) dephasing of each electron spin; the FMN• and FMNH• hyperfine sets; field responses evaluated at k_STD_ = 3 × 10^7^ or 10^6^ s^−1^; and dephasing contrasts of 10^6^ versus 10^8^ s^−1^ or 3 × 10^7^ s^−1^. These spin calculations used a 0.05 Å distance grid with 1,500 Schulten–Wolynes samples per point. In the re-implemented spin models, the grid position of the single highest score depended on sampling seed, bin width and smoothing (range 5.5–6.0 Å in 5XTD), so top-decile statistics are reported as the primary measure.

### Exchange–hyperfine balance and singlet–triplet mixing

The distance-dependent exchange interaction was compared with the effective FMN hyperfine interaction to determine whether the independently identified hotspot coincided with a regime of competing spin interactions.(*11*) Exchange–hyperfine balance was evaluated as a mechanistic diagnostic describing the degree to which the magnitudes of J(r) and a_HF_ approached one another. The balance metric was greatest when exchange and effective hyperfine interactions were comparable and decreased when either interaction strongly dominated.

The singlet–triplet mixing-amplitude proxy was the maximum time-domain departure of the singlet population from its initial value, averaged over the Schulten–Wolynes draws under the low-dephasing condition. It is an amplitude measure, not a transition rate or a measured mixing efficiency. Hyperfine/Zeeman evolution dominates this quantity in the present parameter set. The mixing proxy and exchange–hyperfine balance were calculated for interpretation only and were excluded from the composite score.

### Structure-informed perturbation analysis

The effect of local structural variability on the modelled singlet reaction yield was evaluated using stochastic perturbations of the FMN–O_2_ separation coordinate. The distance-dependent singlet-yield profile generated in the Figure 2 analysis was sorted according to FMN–O_2_ separation and interpolated using a shape-preserving piecewise cubic Hermite interpolating polynomial. This interpolation generated a continuous singlet-yield function, Y _s_(r), while minimising artificial oscillations between adjacent distance-grid values.

The primary singlet-yield profile used for the perturbation analysis was the low-dephasing geomagnetic-field yield when available. The hotspot interval was obtained from the top-10% hotspot bins defined in the preceding structural analysis. When the independently generated hotspot-bin dataset was unavailable, the interval was derived from distance-grid points at or above the 90th percentile of the Figure 2 composite hotspot score.

The selected hotspot bins were converted into a continuous interval by extending the lower and upper boundaries by one-half of the median distance-grid spacing. Because the corrected hotspot lies beyond 5 Å, the long-distance control region used previously (beginning at the greater of 6.5 Å or 1.0 Å beyond the upper hotspot boundary) would overlap the hotspot in 5XTD and lie outside the modelled domain in 9TI4. Hotspot configurations were therefore compared with all other accessible baseline configurations.

### Stochastic structural-fluctuation simulations

A total of 20,000 baseline FMN–O_2_ distances were sampled with replacement from the structure-derived accessible oxygen ensemble. When the individual accessible-configuration dataset was unavailable, baseline distances were sampled from the Figure 2 distance grid with probabilities proportional to the normalised structure-derived accessibility profile. Each baseline configuration was subjected to 128 independent stochastic perturbations at each of five structural-fluctuation amplitudes:

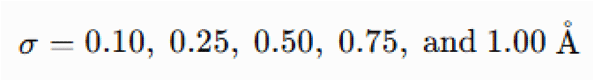

For a baseline separation r_0_, perturbed distances were generated according to

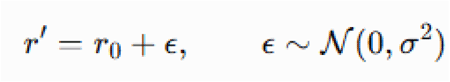

Perturbed distances falling outside the modelled 2.5–9.0 Å domain were constrained to the nearest domain boundary:

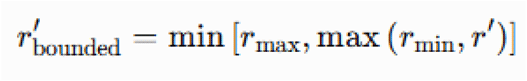

Thus, the numerical implementation used boundary clipping rather than rejection-based truncated-Gaussian resampling. The singlet-yield proxy for each perturbed configuration was obtained from the shape-preserving interpolated yield function:

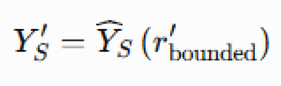

For each baseline configuration and fluctuation amplitude, the mean, variance, and standard deviation of the 128 perturbed singlet-yield values were calculated. Reaction-yield heterogeneity was quantified as

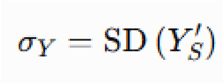

The baseline configuration, rather than each individual perturbation, was treated as the unit of analysis for hotspot–non-hotspot comparisons.

### Structural-response amplification

Structural-response amplification was quantified for each baseline configuration as the ratio of singlet-yield variance to the realised variance of the perturbed FMN–O_2_ separation:

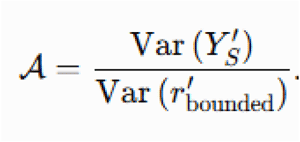

This quantity describes the amount of modelled singlet-yield variance generated per unit of structural variance. Amplification values were subsequently grouped into 0.10-Å baseline-distance bins to construct the distance-dependent amplification landscape.

For each distance bin and fluctuation amplitude, the mean amplification factor and standard error of the mean were calculated. Distance bins containing fewer than 10 baseline configurations were excluded from the binned amplification analysis. Because variance ratios may become sensitive to small denominator values, amplification profiles were interpreted together with the absolute singlet-yield heterogeneity and the underlying distance dependence of the unperturbed singlet-yield profile.

### Statistical analysis

Differences in singlet-yield heterogeneity between hotspot and non-hotspot baseline configurations were evaluated independently for each structural-fluctuation amplitude using a two-sided Mann–Whitney U test. For each fluctuation amplitude, the mean singlet-yield standard deviation was calculated separately for hotspot and non-hotspot baseline configurations. The hotspot-to-non-hotspot heterogeneity ratio was calculated as

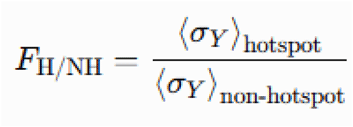

For graphical presentation, uncertainty around the mean heterogeneity was represented as the approximate 95% confidence interval calculated as

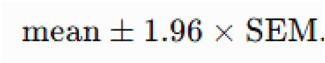

The baseline FMN–O_2_ configuration was treated as the statistical unit. Individual stochastic perturbations generated from the same baseline configuration were used to estimate within-configuration yield variability and were not treated as independent observations in the hotspot–non-hotspot statistical comparison. All statistical tests were two-sided. Exact sample sizes and P values are reported in the corresponding figure or source-data files.

## Software and reproducibility

Structural coordinate processing used Biopython; numerical analysis and plotting used Python, NumPy, pandas and SciPy. The piecewise cubic Hermite interpolating polynomial was implemented with SciPy. Spin propagation used custom Python/NumPy code with hyperfine constants from RadicalPy. The pseudo-random number generator was initialised with the stated fixed seed, 260705, for reproducibility of the stochastic calculations.

## Declarations

### Ethics approval and consent to participate

N/A

### Consent for publication

N/A

### Availability of data and materials

The analysis codes and datasets used and/or generated during the current study are available from Dr. Ji-Yong Sung upon reasonable request. Interested researchers may contact Dr. Sung via email at to obtain access to the relevant materials.

### AI Use Declaration

AI tools assisted with language revision and checks for repetition and internal consistency.

### Authors’ contributions

Conceptualization & Investigation: JYS, JHC; Methodology: JYS; Data analysis: JYS; Writing-original draft: JYS; Writing-review & editing: JYS, JHC; Supervision: JYS, JHC; Project administration: JYS, JHC; Funding acquisition: JHC; Interpretation of the results: JYS, LMA, JHC. All authors have read and agreed to the published version of the manuscript.

## Acknowledgements

N/A

## Supplementary Information

### Supplementary Note 1: Exchange and point-dipole coupling, extended to 2.5–20 Å

We added an axial point-dipole term to the exchange Hamiltonian, using D(r) = μ g²μB²/(4πhr³) and the operator D(r)[S1·S2 − 3S1zS2z]; the previous version used the opposite overall sign. The FMN–O_2_ separation vector was assumed to be collinear with the field axis. This fixed orientation gives a strong dipolar contribution but does not represent the angular average of the anisotropic interaction. D(r) falls as r ³ and therefore decreases more slowly than the exponential exchange term. In this calculation, D(r) is approximately 3.6 mT at 8 Å and 0.23 mT at 20 Å, compared with an effective FMN hyperfine field of 1.23 mT. Near 2.5–4 Å, the calculated dipolar coupling is about 40–120 mT. Such large electron– electron interactions can suppress magnetic-field sensitivity, as discussed for radical-pair reactions by Efimova and Hore (2008). The collinear calculation tests the influence of a substantial dipolar interaction on the composite score; it does not establish the coupling experienced by an FMN–superoxide pair in complex I.

With FMN-overlapping positions excluded, the 1,300,000-candidate 2.5–20 Å analysis of 5XTD gave no localized composite-score maximum with exchange alone (highest grid value at the 20 Å boundary) and a maximum at 6.05 Å with exchange plus dipolar coupling (top-decile mean 8.18 Å, SD 1.43 Å). The corresponding results for 9TI4 were a boundary maximum and a maximum at 7.85 Å (mean 9.53 Å, SD 1.82 Å) (Table 1). The near-contact maxima at 2.75–3.35 Å and the local features at 4.65–5.35 Å reported in the previous version were obtained with a steric test that omitted FMN atoms and are not reproduced. The dipolar calculation also assumes a collinear geometry; orientation-dependent effects remain untested.

**Figure S1.**
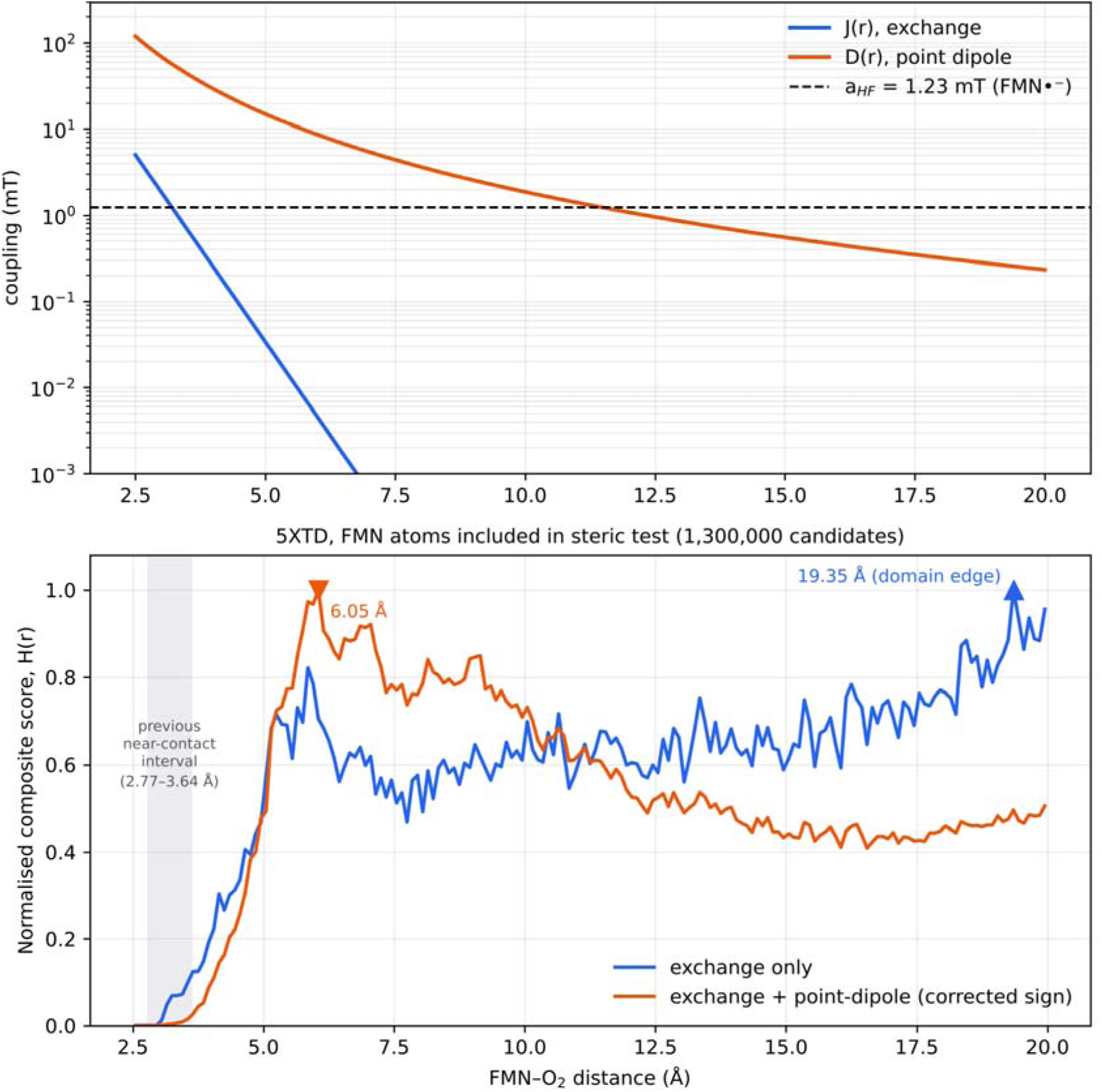
Point-dipole and extended-domain calculations. Top: exchange J(r), collinear point-dipole coupling D(r) and reference FMN hyperfine field over separation. Bottom: composite scores over 2.5–20 Å for exchange alone or exchange plus point-dipole coupling, recalculated for 5XTD with FMN atoms included in the steric test (1,300,000 candidates), the corrected dipolar sign and the re-implemented spin model. The exchange-only score has no localized maximum (highest grid value at the 20 Å edge); with dipolar coupling the maximum lies at 6.05 Å. Grey band, near-contact interval of the previous version.

The raw field-response curves from the initial 2.5–9.0 Å sweep are shown in Fig. S2. Their marked maxima indicate the largest sampled values within that restricted range. The longer-range, higher-precision calculation in Fig. S4 shows that these markers do not identify isolated physical peaks.

**Figure S2.**
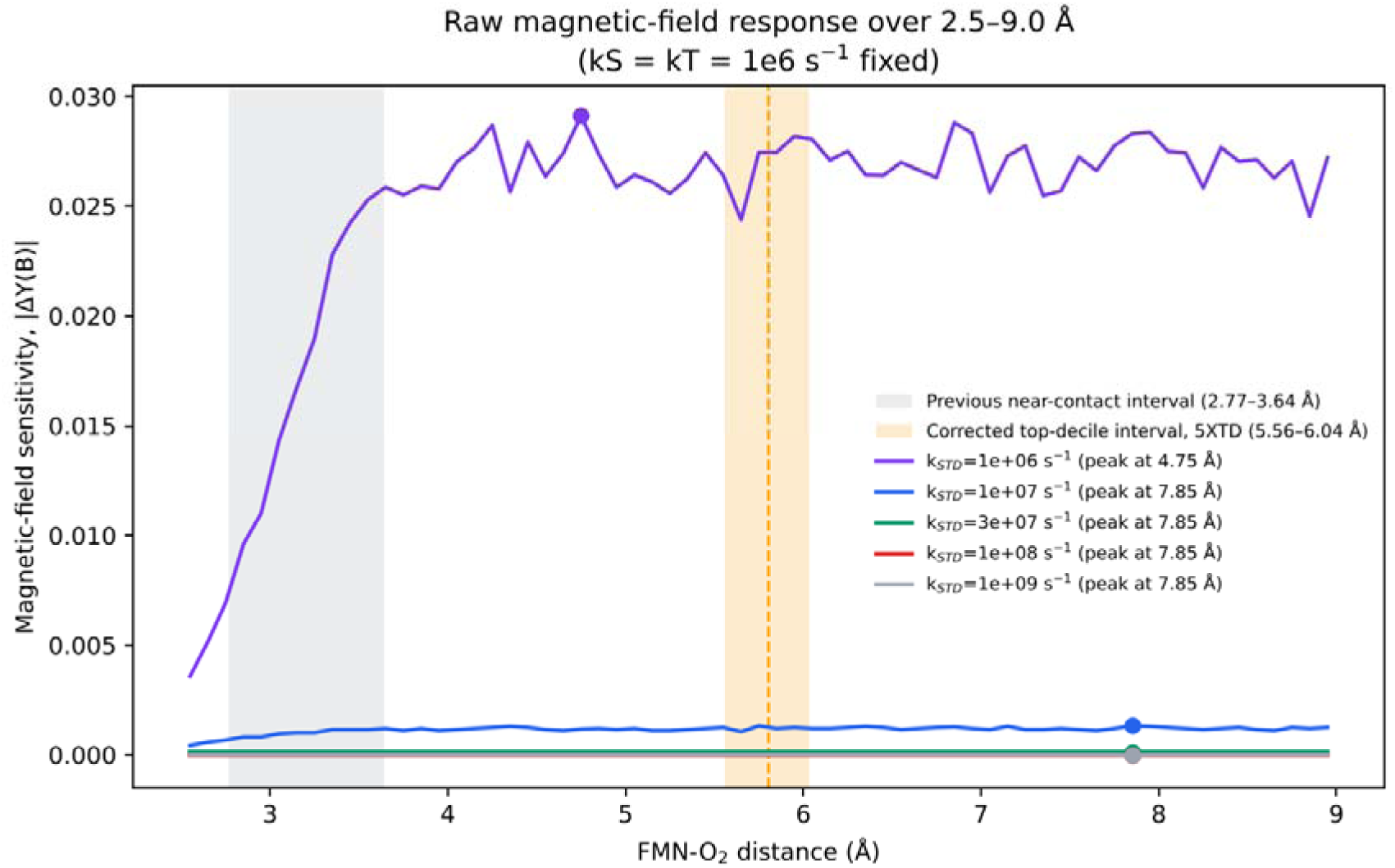
Raw magnetic-field response over the original 2.5–9.0 Å interval. Curves show |ΔY(B)| for k_STD_ = 10 –10□s□¹ at k_S_ = k_T_ = 10□s□¹. The marked 4.75 and 7.85 Å positions are the largest sampled bins in that restricted window; the 4.75 Å position lies below, and the 7.85 Å position above, the corrected 5XTD percentile interval (5.56–6.04 Å). The orange band shows the corrected 5XTD top-decile interval (dashed line, mean 5.80 Å) and the grey band the near-contact interval of the previous version. The extended calculation in Fig. S4 shows that they do not define narrow field-response peaks.

**Figure S3.**
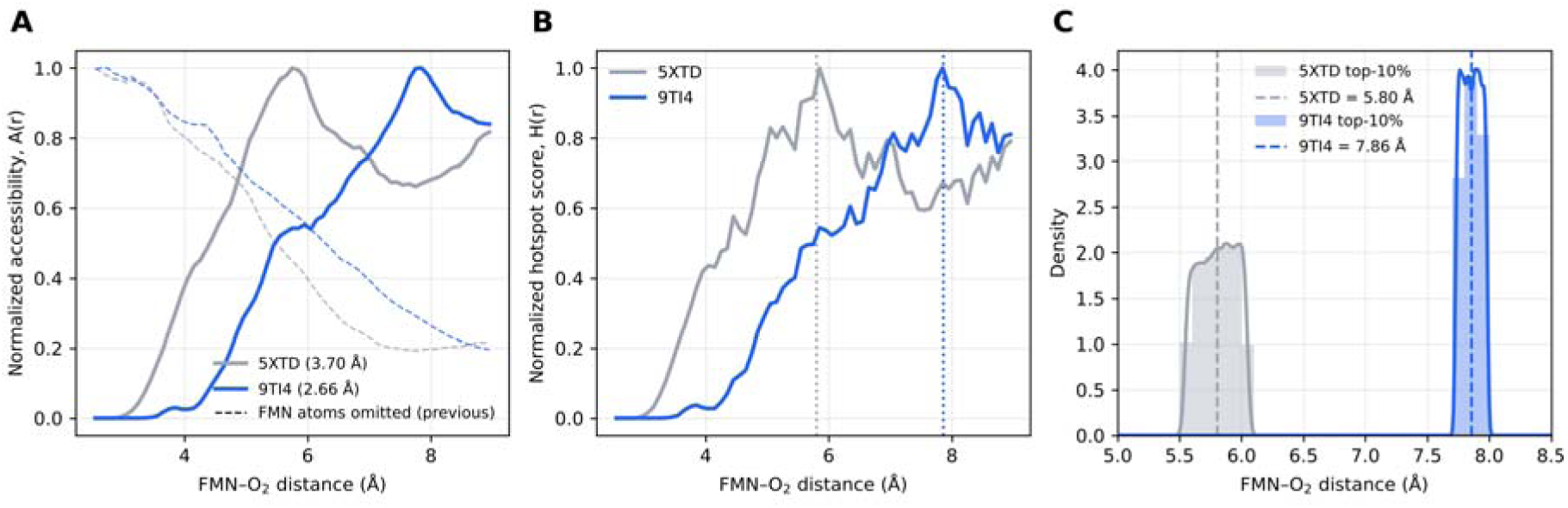
Structural-input comparison. The 5XTD structure (3.70 Å resolution) and independently solved 9TI4 structure (2.66 Å resolution) were analysed with the same steric filter, including FMN atoms, and spin parameters. (A) Normalised oxygen acceptance fraction A(r); dashed lines, previous version with FMN atoms omitted. (B) Composite score H(r); dotted lines, top-decile means. (C) Distributions of top-decile distances. Top-decile means were 5.80 Å (5XTD) and 7.86 Å (9TI4), with 2.5th–97.5th percentile intervals of 5.56–6.04 and 7.74–7.98 Å, respectively.

### Supplementary Note 2: Confirmatory 1,300,000-candidate re-analysis and high-precision |ΔY(B)| plateau check (12 Aug 2026)

We investigated the broad distance distributions obtained after extending the sampling domain to 20 Å. The accessibility calculation was repeated with 1,300,000 trial centroids using the same steric filter, and raw |ΔY(B)|(r) was recalculated in the re-implemented spin model with 1,500 Schulten–Wolynes samples per distance using common random numbers at both magnetic fields. The tests distinguished finite-sampling effects from the influence of the long-range field-response profile.

With FMN atoms included in the steric test, the larger structural sample had an accessible fraction of 7.6% for 5XTD (7.5% for 9TI4; 8.4% when FMN atoms were omitted, as in the previous version). It produced no near-contact score maximum: with exchange alone, the score formed a broad plateau beyond about 5 Å, and with dipolar coupling the maximum lay at 6.05 Å (5XTD) or 7.85 Å (9TI4) (Table 1). The broad extended-range distributions therefore do not result from limited sampling density.

With improved spin sampling in the re-implemented spin model, with the corrected dipolar sign, exchange-only |ΔY(B)| was suppressed near contact at k_STD_ ≤ 3 × 10^7^ s^−1^ and approached a nearly constant value by about 5 Å (Fig. S4). Including the collinear dipolar term delayed the approach to the plateau to approximately 8–10 Å and produced a broad maximum between about 6 and 13 Å, which moved to shorter separation as k_STD_ increased, followed by a slow decline; a narrow feature at 2.75 Å lies inside the region excluded by FMN. None of these curves showed a narrow maximum in the accessible range. The 4.75 and 7.85 Å markers in Fig. S2 reflect selection of individual sampled bins in the earlier, shorter calculation; they should not be assigned mechanistic significance.

We multiplied the raw field-response curve by normalised accessibility A(r) to examine where the model places geometrically weighted field response over 2.5–20 Å. With the corrected accessibility, the integrated share for the 4.75–7.85 Å interval was 19% in 5XTD and 13% in 9TI4 with exchange alone, and 4–22% with exchange plus dipolar coupling, depending on k_STD_. Within the 2.77–3.64 Å interval it was at most 0.2%, because almost no accessible positions remain there (previous version, 0.1–12%). These fractions depend on interval width, steric weighting and the spin Hamiltonian; they are not fractions of measured ROS production

The extended calculations separate the location of a composite-score peak from the breadth of high-scoring configurations and from raw magnetic-field sensitivity. After correction of the steric test, the composite-score maximum is not near contact, and neither the larger accessible ensemble nor the more precise field-response calculation supports a narrow second magnetic-response optimum at larger separation.

**Figure S4.**
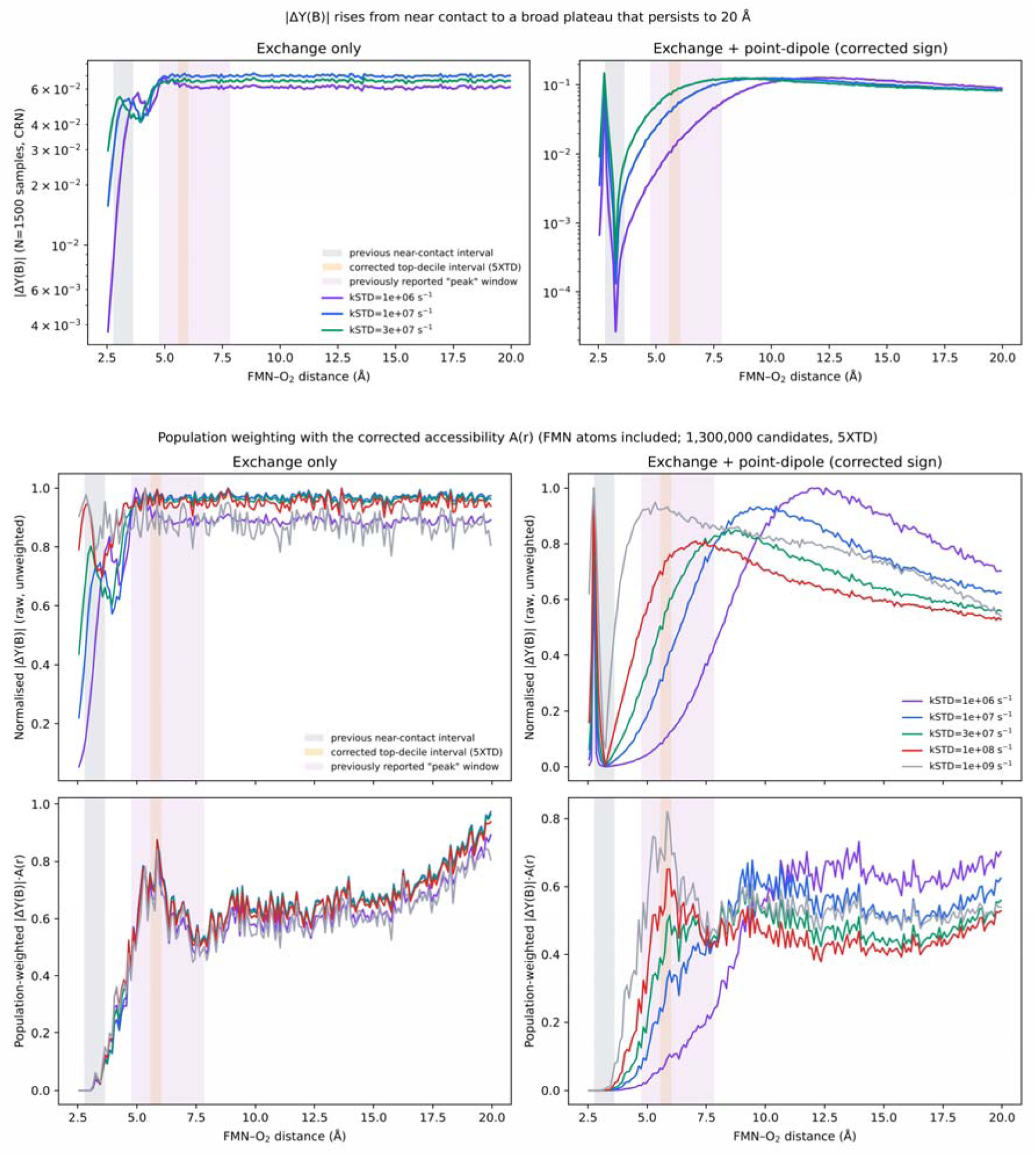
Higher-precision field response across 2.5–20 Å (re-implemented spin model; 1,500 Schulten–Wolynes samples per distance with common random numbers). Top: raw |ΔY(B)| for k_STD_ = 10^6^, 10^7^ and 3 × 10^7^ s^−1^, using exchange alone or exchange with a collinear point-dipole term (corrected sign). Bottom: normalised raw |ΔY(B)| (upper row) and accessibility-weighted response |ΔY(B)|·A(r) (lower row) for k_STD_ = 10^6^–10^9^ s^−1^, using the corrected accessibility of 5XTD (FMN atoms included; 1,300,000 candidates). Grey band, near-contact interval of the previous version (2.77–3.64 Å); orange band, corrected 5XTD top-decile interval (5.56–6.04 Å); purple band, 4.75–7.85 Å range of the largest sampled bins in Fig. S2. The raw responses approach broad plateaus without a narrow maximum in the accessible range; the narrow feature at 2.75 Å in the dipolar model lies inside the region excluded by FMN, and accessibility weighting removes all contribution below about 3.5 Å.

**Figure S5.**
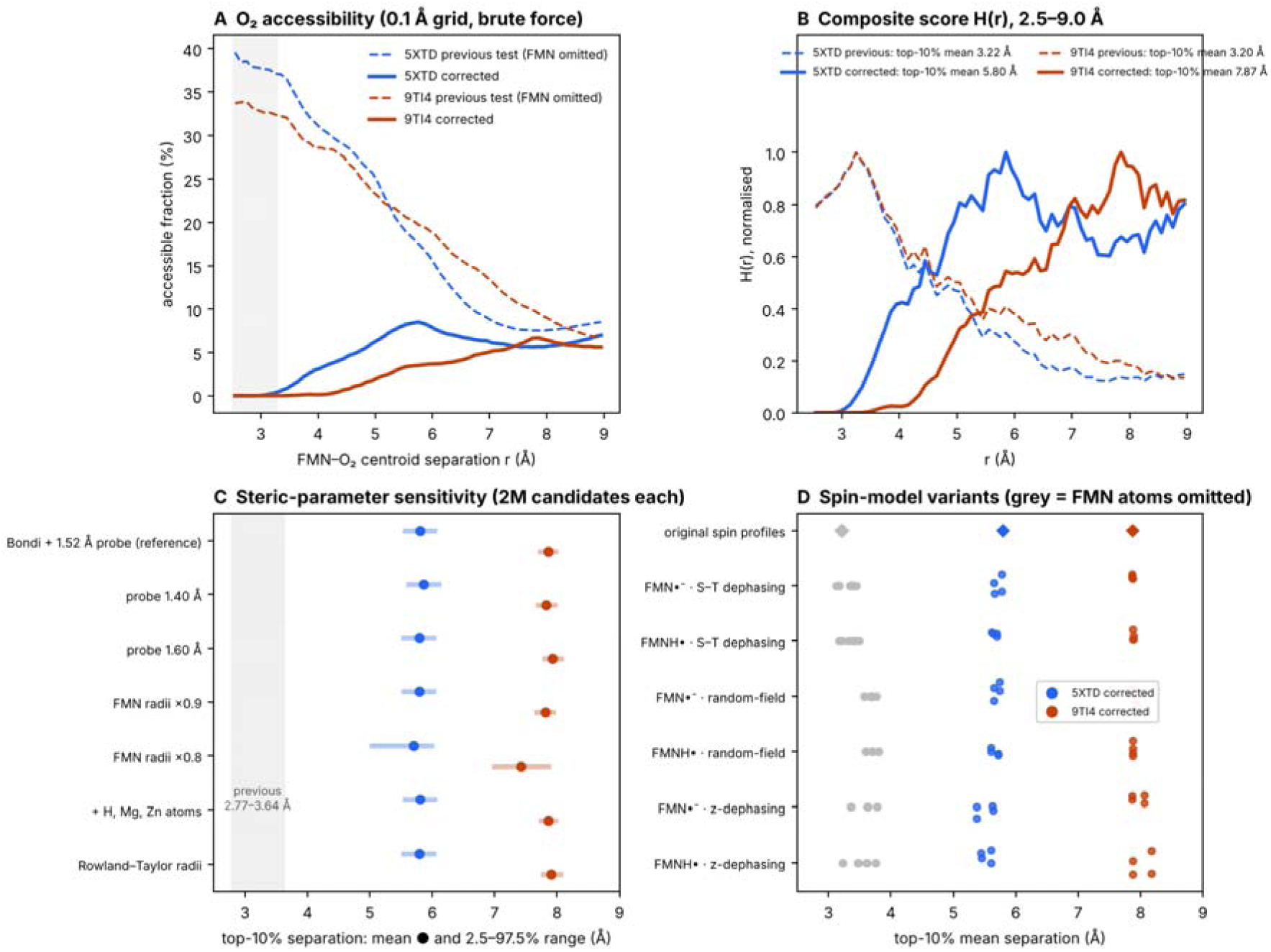
Correction of the steric test in two complex I structures. (A) Accessible fraction of candidate O_2_ centroids on a regular 0.1 Å grid, calculated by brute-force distance evaluation, with FMN atoms omitted (dashed; previous version) or included (solid). (B) Composite score H(r) over 2.5–9.0 Å, using the original spin-response profiles, for 5XTD (blue) and 9TI4 (orange), with FMN atoms omitted (dashed) or included (solid). (C) Top-decile mean separations (dots) and 2.5th–97.5th percentile ranges (bars) for steric-parameter variants, each with 2,000,000 candidates. Grey band, near-contact interval reported in the previous version. (D) Top-decile mean separations with the original spin-response profiles (diamonds) and for re-implemented spin-model variants (three dephasing models × two flavin hyperfine sets × four response-metric definitions); grey, FMN atoms omitted.

## References

1. Gerald S. Shadel, Tamas L. Horvath, Mitochondrial ROS Signaling in Organismal Homeostasis. Cell 163, 560–569 (2015).

2. Laura A. Sena, Navdeep S. Chandel, Physiological Roles of Mitochondrial Reactive Oxygen Species. Molecular Cell 48, 158–167 (2012).

3. Michael P. Murphy, How mitochondria produce reactive oxygen species. Biochemical Journal 417, 1–13 (2008).

4. J. Y. Sung, J. H. Cheong, Quantum biology: From mechanisms to medicine. Clinical and Translational Medicine 16, (2026).

5. J. Y. Sung, J. H. Cheong, Quantum medicine: A quantum–mechanical framework for redox biology, disease and precision medicine. Clinical and Translational Medicine 16, (2026).

6. J.-Y. Sung, J.-H. Cheong, Quantum thermodynamics with information and coherence. Quantum Information Processing 25, (2026).

7. J. Langowski, R. J. Usselman, I. Hill, D. J. Singel, C. F. Martino, Spin Biochemistry Modulates Reactive Oxygen Species (ROS) Production by Radio Frequency Magnetic Fields. PLoS ONE 9, (2014).

8. C. C. Moser, S. E. Chobot, C. C. Page, P. L. Dutton, Distance metrics for heme protein electron tunneling. Biochimica et Biophysica Acta (BBA) - Bioenergetics 1777, 1032–1037 (2008).

9. C. C. Page, C. C. Moser, P. L. Dutton, Mechanism for electron transfer within and between proteins. Current Opinion in Chemical Biology 7, 551–556 (2003).

10. N. Lambert et al., Quantum biology. Nature Physics 9, 10–18 (2012).

11. H. Zadeh-Haghighi, C. Simon, Magnetic field effects in biology from the perspective of the radical pair mechanism. Journal of The Royal Society Interface 19, (2022).

12. R. J. Usselman et al., The Quantum Biology of Reactive Oxygen Species Partitioning Impacts Cellular Bioenergetics. Scientific Reports 6, (2016).

13. J. Hirst, Mitochondrial Complex I. Annual Review of Biochemistry 82, 551–575 (2013).

14. L. Kussmaul, J. Hirst, The mechanism of superoxide production by NADH:ubiquinone oxidoreductase (complex I) from bovine heart mitochondria. Proceedings of the National Academy of Sciences 103, 7607–7612 (2006).

15. T. P. Fay, L. P. Lindoy, D. E. Manolopoulos, P. J. Hore, How quantum is radical pair magnetoreception? Faraday Discussions 221, 77–91 (2020).

16. M. J. Golesworthy et al., Singlet–triplet dephasing in radical pairs in avian cryptochromes leads to time-dependent magnetic field effects. The Journal of Chemical Physics 159, (2023).

17. L. M. Antill, E. Vatai, RadicalPy: A Tool for Spin Dynamics Simulations. Journal of Chemical Theory and Computation 20, 9488–9499 (2024).

18. P. J. Hore, H. Mouritsen, The Radical-Pair Mechanism of Magnetoreception. Annual Review of Biophysics 45, 299–344 (2016).

19. M. C. J. Denton et al., Magnetosensitivity of tightly bound radical pairs in cryptochrome is enabled by the quantum Zeno effect. Nature Communications 15, (2024).

20. V. Bezchastnov, T. Domratcheva, The arrangement of anisotropic spin couplings can optimize sensitivity of the cryptochrome radical pair to the direction of geomagnetic field. Scientific Reports 16, (2026).

21. D. Khurana et al., Sensing of magnetic field effects in radical-pair reactions using a quantum sensor. Physical Review Research 6, (2024).

22. M. D. Nguyen et al., Structural basis for late maturation steps of mitochondrial respiratory chain complex IV within the human respirasome. Nature Communications 17, (2026).

23. D. Esterházy, M. S. King, G. Yakovlev, J. Hirst, Production of Reactive Oxygen Species by Complex I (NADH:Ubiquinone Oxidoreductase) from Escherichia coli and Comparison to the Enzyme from Mitochondria. Biochemistry 47, 3964–3971 (2008).

24. Y. Jiang, S. Kirmizialtin, I. C. Sanchez, Dynamic void distribution in myoglobin and five mutants. Scientific Reports 4, (2014).

25. R. Kitahara, Y. Yoshimura, M. Xue, T. Kameda, F. A. A. Mulder, Detecting O2 binding sites in protein cavities. Scientific Reports 6, (2016).

26. A. Bondi, van der Waals Volumes and Radii. The Journal of Physical Chemistry 68, 441–451 (2002).

27. I. Polyakov, A. Kulakova, A. Nemukhin, Computational Modeling of the Interaction of Molecular Oxygen with the miniSOG Protein—A Light Induced Source of Singlet Oxygen. Biophysica 3, 252–262 (2023).

28. J. Deng, Q. Cui, Efficient Sampling of Cavity Hydration in Proteins with Nonequilibrium Grand Canonical Monte Carlo and Polarizable Force Fields. Journal of Chemical Theory and Computation 20, 1897–1911 (2024).

29. C.-L. Teng, R. G. Bryant, Mapping Oxygen Accessibility to Ribonuclease A Using High-Resolution NMR Relaxation Spectroscopy. Biophysical Journal 86, 1713–1725 (2004).

30. M. S. Al-Abdul-Wahid et al., A Combined NMR and Molecular Dynamics Study of the Transmembrane Solubility and Diffusion Rate Profile of Dioxygen in Lipid Bilayers. Biochemistry 45, 10719–10728 (2006).

31. C.-L. Teng, B. Hinderliter, R. G. Bryant, Oxygen Accessibility to Ribonuclease A: Quantitative Interpretation of Nuclear Spin Relaxation Induced by a Freely Diffusing Paramagnet. The Journal of Physical Chemistry A 110, 580–588 (2005).

32. G. J. Pažėra, T. P. Fay, I. A. Solov’yov, P. J. Hore, L. Gerhards, Spin Dynamics of Radical Pairs Using the Stochastic Schrödinger Equation in MolSpin. Journal of Chemical Theory and Computation 20, 8412–8421 (2024).

33. C. K. Austvold, S. M. Keable, M. Procopio, R. J. Usselman, Quantitative measurements of reactive oxygen species partitioning in electron transfer flavoenzyme magnetic field sensing. Frontiers in Physiology 15, (2024).

34. O. Efimova, P. J. Hore, Role of Exchange and Dipolar Interactions in the Radical Pair Model of the Avian Magnetic Compass. Biophysical Journal 94, 1565–1574 (2008).

35. I. A. Solov’yov, D. E. Chandler, K. Schulten, Magnetic Field Effects in Arabidopsis thaliana Cryptochrome-1. Biophysical Journal 92, 2711–2726 (2007).

36. C. Riplinger et al., Interaction of Radical Pairs Through-Bond and Through-Space: Scope and Limitations of the Point−Dipole Approximation in Electron Paramagnetic Resonance Spectroscopy. Journal of the American Chemical Society 131, 10092–10106 (2009).

37. P. J. Hore, K. L. Ivanov, M. R. Wasielewski, Spin chemistry. The Journal of Chemical Physics 152, (2020).

38. G. Lindblad, On the generators of quantum dynamical semigroups. Communications in Mathematical Physics 48, 119–130 (1976).

39. R. Haberkorn, Density matrix description of spin-selective radical pair reactions. Molecular Physics 32, 1491–1493 (2006).

40. T. C. Player, P. J. Hore, Viability of superoxide-containing radical pairs as magnetoreceptors. The Journal of Chemical Physics 151, 225101 (2019). 10.1063/1.5129608.

41. J. Deviers, F. Cailliez, A. de la Lande, D. R. Kattnig, Avian cryptochrome 4 binds superoxide. Computational and Structural Biotechnology Journal 26, 11–21 (2024). 10.1016/j.csbj.2023.12.009.

42. K. M. Salerno, J. Domenico, N. Q. Le, C. D. Stiles, I. A. Solov’yov, C. F. Martino, Long-Time Oxygen Localization in Electron Transfer Flavoprotein. Journal of Chemical Information and Modeling 62, 4191–4199 (2022). 10.1021/acs.jcim.2c00430.

43. K. M. Salerno, J. Domenico, N. Q. Le, K. Balakrishnan, R. J. McQuillen, C. D. Stiles, I. A. Solov’yov, C. F. Martino, Long-Time Oxygen and Superoxide Localization in Arabidopsis thaliana Cryptochrome. Journal of Chemical Information and Modeling 63, 6756–6767 (2023). 10.1021/acs.jcim.3c00325.

