## Supplementary figures and images for "Structure-constrained spin responses of an FMN–superoxide radical-pair model in mitochondrial complex I"

### Supplementary Figure1

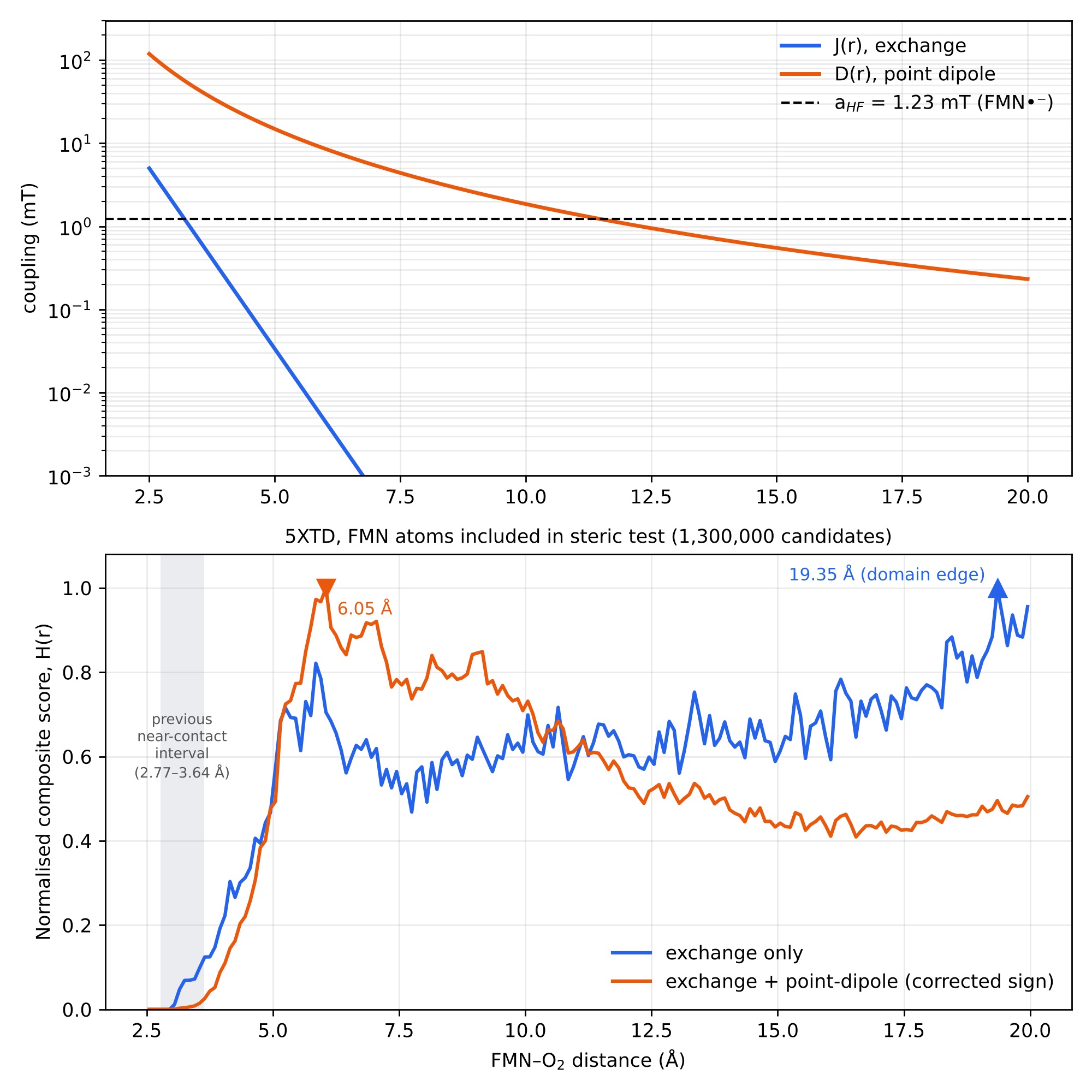

### Supplementary Figure2

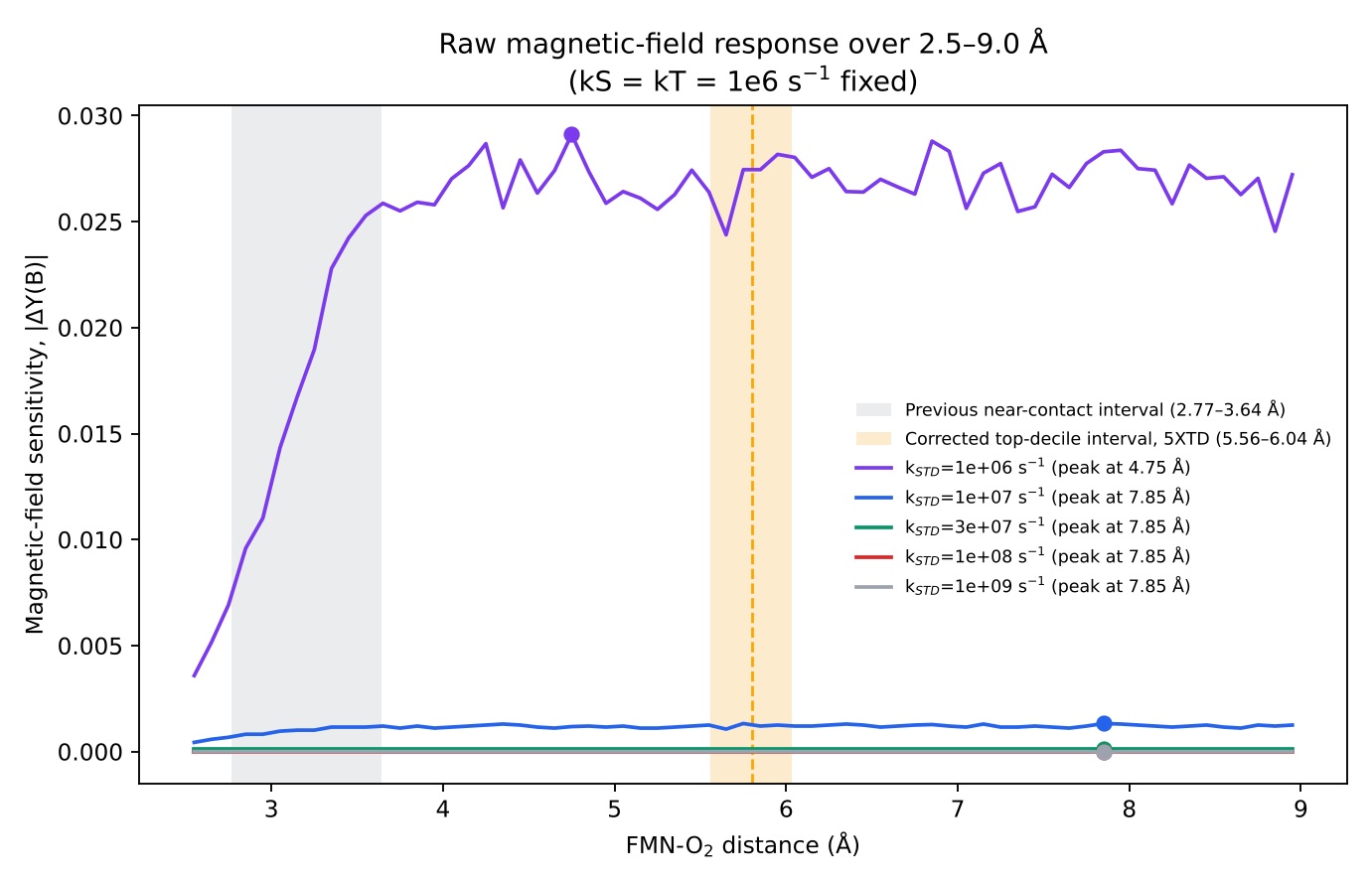

### Supplementary Figure3

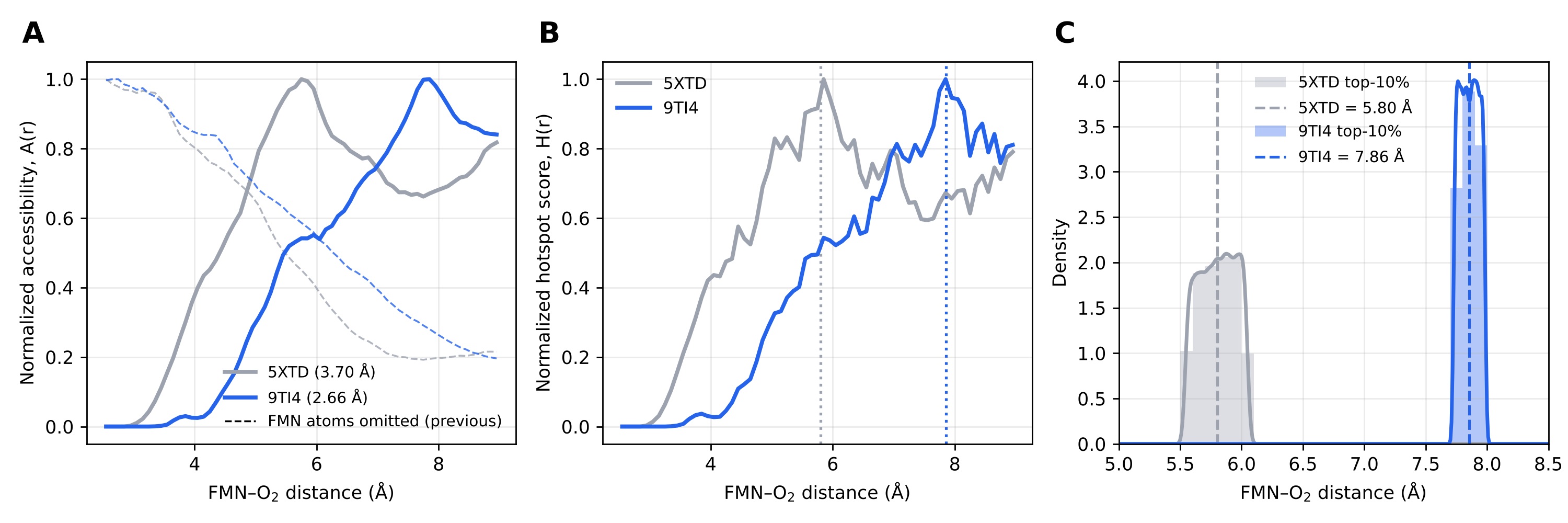

### Supplementary Figure4_1

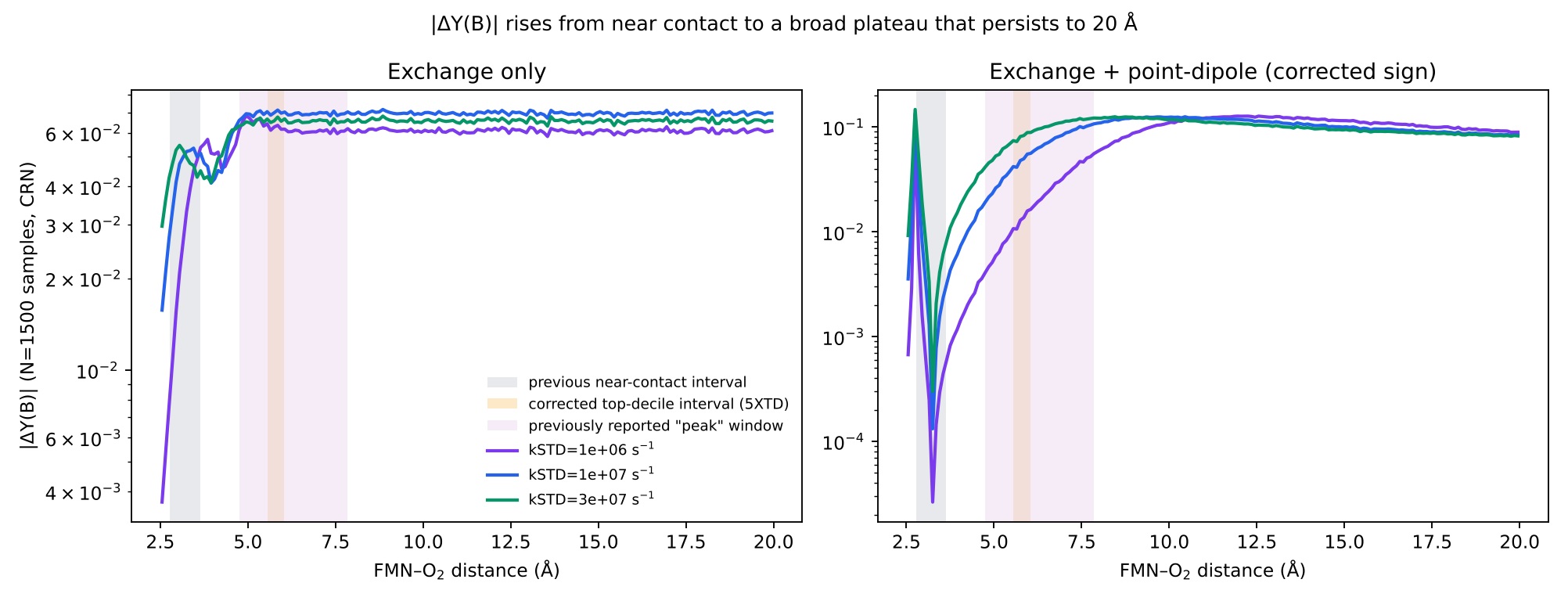

### Supplementary Figure4_2

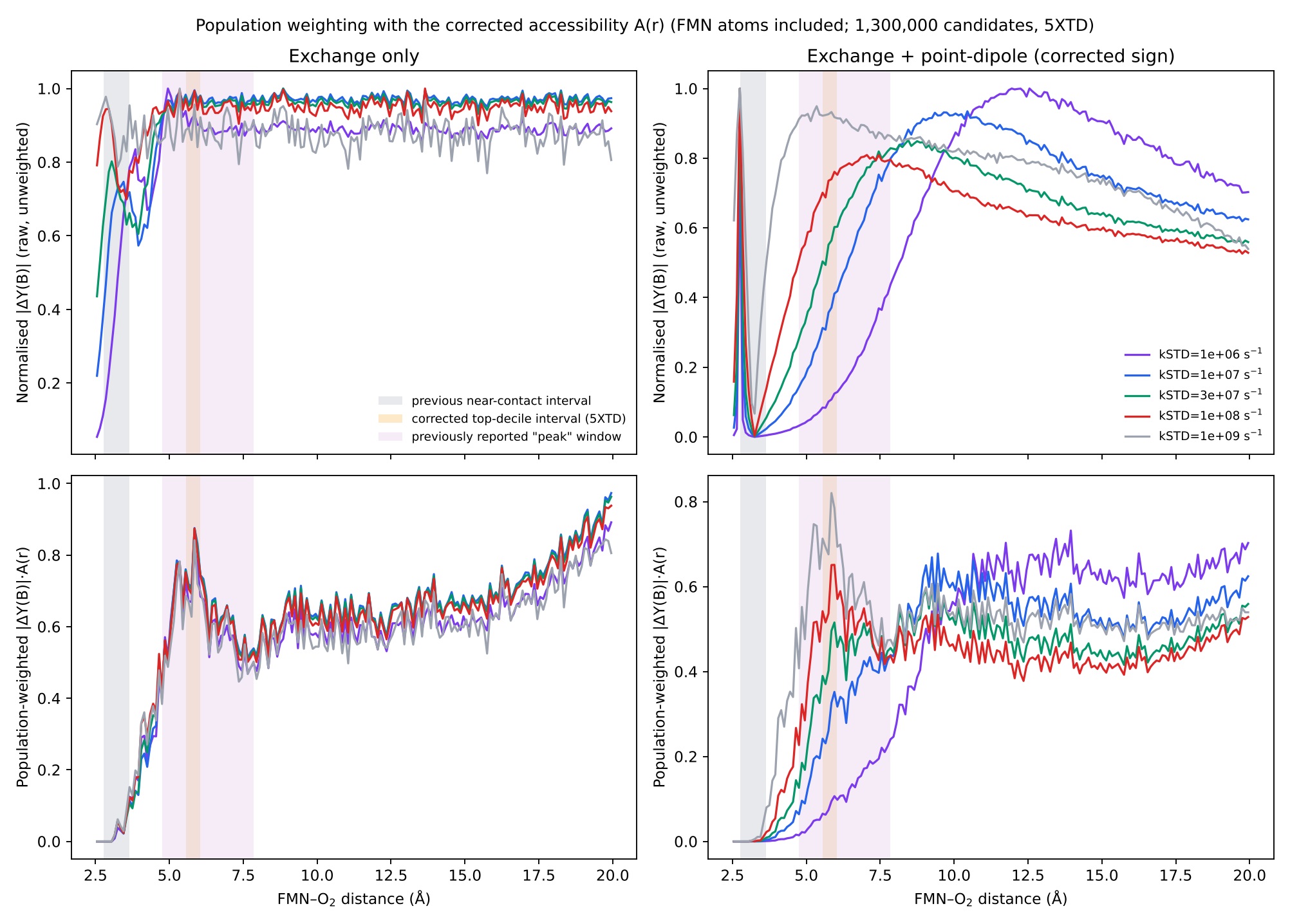

### Supplementary Figure5

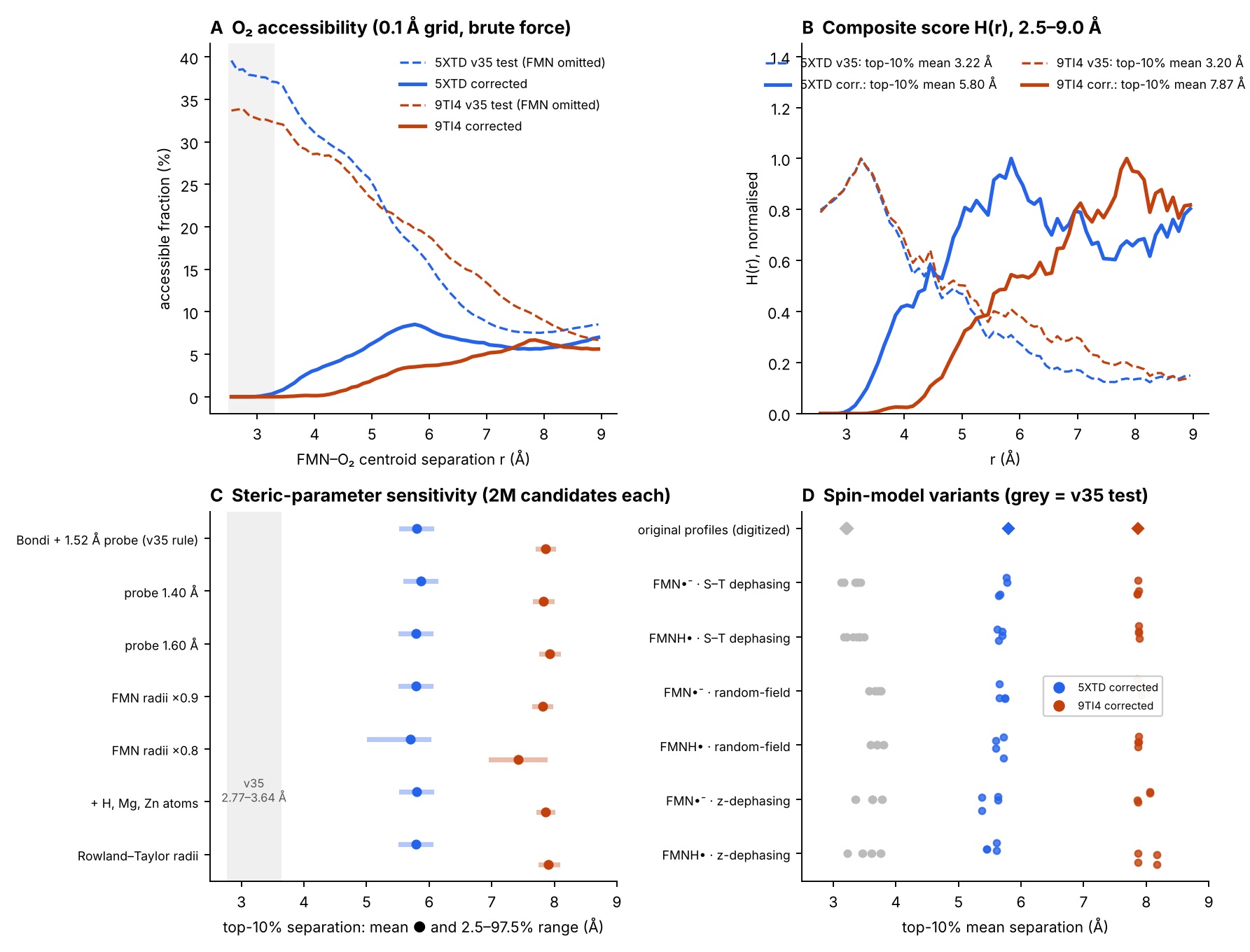
