## Supplementary Information for "Structure-constrained spin responses of an FMN–superoxide radical-pair model in mitochondrial complex I"

**Supplementary Note 1: Exchange and point-dipole coupling, extended to 2.5–20 Å**

A point-dipole electron–electron coupling term, D(r), was computed from first principles (D(r) = μ_0_g^2^μ_B_^2^/4πhr^3^) and added to the spin Hamiltonian as an axial term, −(3S_1z_S_2z_ − S_1_·S_2_), on top of the existing exchange term J(r)S_1_·S_2_. This treats the FMN–O_2_ vector as collinear with the Zeeman quantisation axis. Because the axial dipolar coefficient, D(r)(3cos²θ−1), is maximal in magnitude at θ=0, this collinear geometry corresponds to the orientation that maximises the dipolar term's potential to compete with or suppress hyperfine-driven singlet–triplet mixing — i.e. the case in which D(r) competes most strongly with hyperfine-driven mixing. The overall minus sign corrects the previous implementation, which used the opposite sign of the dipolar term. The collinear treatment remains an explicit simplification of the true anisotropic, orientation-dependent dipolar tensor, adopted here as a first-pass estimate rather than a full orientation-averaged treatment. D(r) decays as r^-3^, much more slowly than the exponentially decaying J(r): by r≈8 Å, J(r) is essentially zero while D(r) (3.6 mT) still exceeds the FMN hyperfine coupling (1.23 mT), and D(r) remains non-negligible (0.23 mT) even at r=20 Å. At near-contact separations (2.5–4 Å), D(r) (40–120 mT) exceeds both J(r) (2–5 mT) and the hyperfine coupling by more than an order of magnitude, consistent with the concern that exchange and dipolar coupling may compete with or suppress hyperfine-driven singlet–triplet mixing in this region. This is consistent with the analysis of Efimova & Hore (2008), who showed for the avian radical-pair compass that both the exchange and dipolar couplings must be small relative to the hyperfine interaction — or must very nearly cancel one another — for the radical-pair mechanism to retain appreciable magnetic-field sensitivity; large, uncancelled exchange or dipolar coupling was shown to quench the anisotropic magnetic response. Our finding that D(r) exceeds aHF throughout the sampled near-contact domain (falling below it only beyond ~11.5 Å) places our near-contact geometry in the regime Efimova and Hore identify as suppressive rather than favourable, and is the direct motivation for reframing the near-contact region as an exchange/dipolar–hyperfine competition regime rather than one favourable for singlet–triplet interconversion. Once positions overlapping FMN are excluded from the accessibility calculation, however, almost no oxygen positions remain in this near-contact region (≤0.2% within 3.3 Å; Supplementary Note 3), so it does not contribute appreciably to the accessibility-weighted score.

The composite hotspot analysis was repeated over an extended 2.5–20 Å domain (requiring a corresponding extension of the structural accessibility ensemble), both without and with the point-dipole term (**Fig. S1**). With positions overlapping FMN excluded from the accessibility ensemble (1,300,000 candidates; Supplementary Note 3), no near-contact maximum appeared in either case. With exchange alone, the score formed a broad plateau beyond about 5 Å and its highest grid value lay at the 20 Å domain boundary in both 5XTD and 9TI4 (top-decile mean 19.6 Å), so no localized maximum could be defined. With the point-dipole term included, the maximum lay at 6.05 Å in 5XTD (top-decile mean 8.18 Å, SD 1.43 Å; 2.5th–97.5th percentiles, 5.39–10.13 Å) and at 7.85 Å in 9TI4 (mean 9.53 Å, SD 1.82 Å; 6.81–13.38 Å). The near-contact maxima reported in the previous version (3.35 Å with exchange alone; 3.25 and 2.75 Å with dipolar coupling, forming an apparent near-bimodal structure) and the smaller local maxima at 4.65–5.35 Å were obtained with a steric test that omitted FMN atoms. Of the positions that test accepted within 3.3 Å of the ring centre, 99–100% overlap FMN atoms, and these features are not reproduced. The dipolar calculations reported here use the corrected sign of the dipolar term, which changed the corrected top-decile statistics by at most 0.05 Å. The collinear point-dipole approximation does not capture orientation-dependent effects that a full anisotropic tensor treatment might reveal.

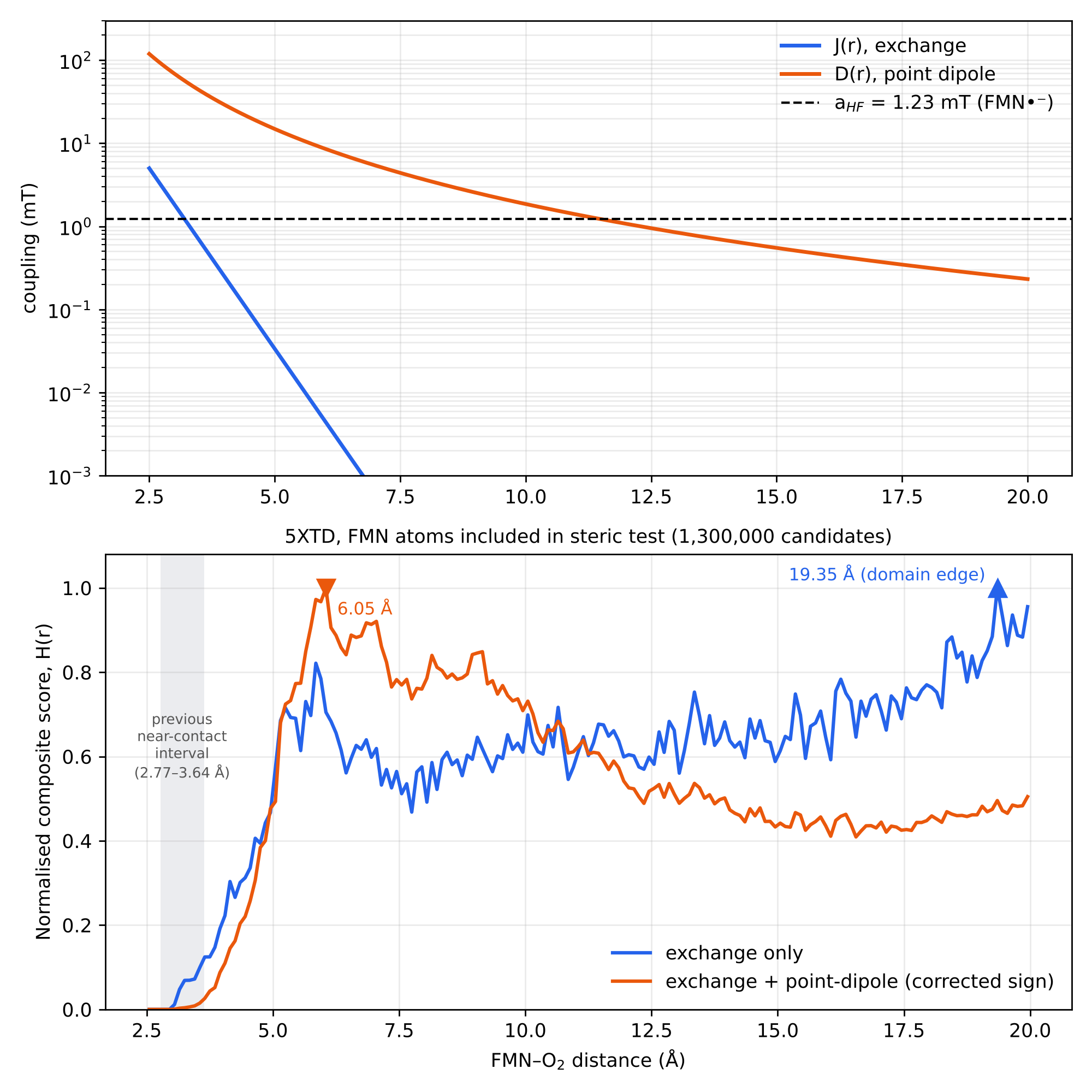

**Figure S1.** Top: exchange coupling J(r) and point-dipole coupling D(r) versus FMN–O_2_ distance (log scale), with the FMN hyperfine coupling shown for reference. Bottom: normalised composite hotspot score, H(r), computed without (exchange only) and with (exchange + point-dipole) the dipolar term, over the extended 2.5–20 Å domain, using the corrected accessibility ensemble (FMN atoms included in the steric test; 1,300,000 candidates, 5XTD), the corrected dipolar sign and a re-implementation of the spin model. The exchange-only score rises to a broad plateau with no localized maximum; the exchange + point-dipole score peaks at 6.05 Å. The near-contact local maxima (2.75, 3.25 and 3.35 Å) shown in the previous version are not reproduced.

As noted in the main text, the raw magnetic-field sensitivity, |ΔY(B)|, does not itself peak at the composite hotspot location. Fig. S2 shows |ΔY(B)| versus FMN–O₂ distance for each k_STD_ value tested in the sweep of Fig. 5B, with the per-curve peak location marked.

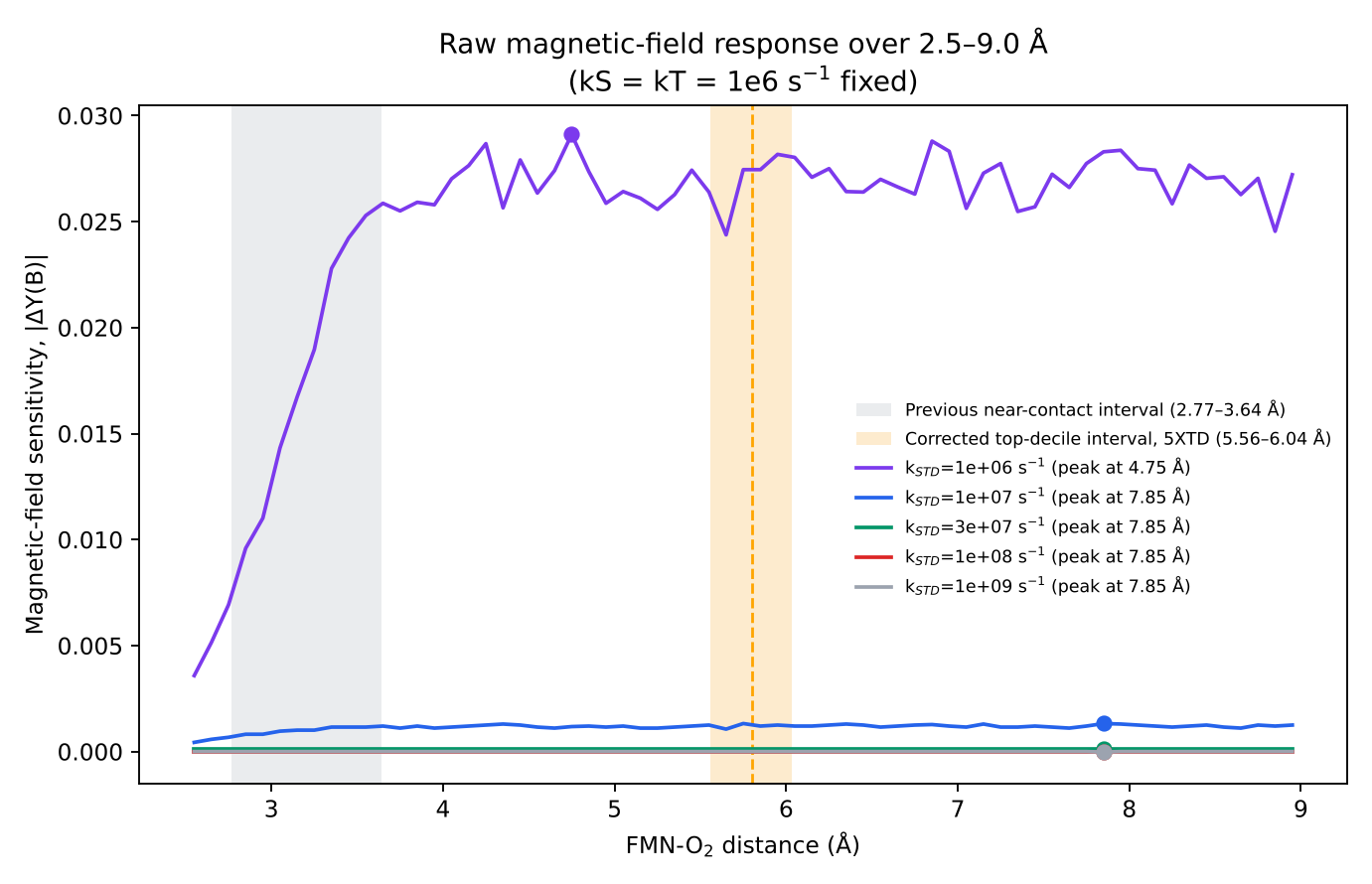

**Figure S2.** The magnetic-field sensitivity, |ΔY(B)|, does not peak at the composite hotspot location. Curves show |ΔY(B)| versus FMN–O₂ distance for k_STD_ = 10^6^–10^9^ s^-1^ (k_S_ = k_T_ = 10^6^ s^-1^ fixed), with per-curve peak markers. The peak location shifts from 4.75 Å (k_STD_=10^6^ s^-1^) to 7.85 Å (all higher k_STD_ values tested), below and above, respectively, the corrected 5XTD top-decile interval (5.56–6.04 Å). The orange band shows the corrected 5XTD top-decile interval (dashed line, mean 5.80 Å); the grey band shows the near-contact interval of the previous version (2.77–3.64 Å).

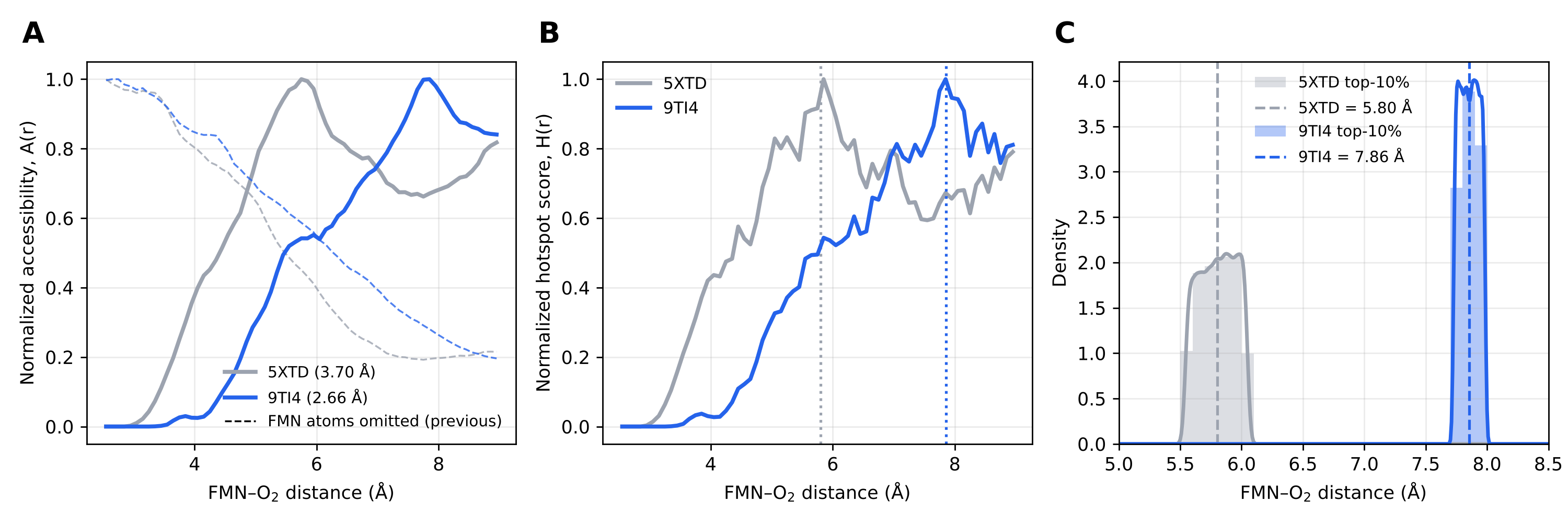

**Figure S3.** Comparison of two complex I structures after correction of the steric test. The accessibility and spin-dynamics pipeline (element-specific van der Waals steric filtering against protein and FMN atoms, followed by Schulten–Wolynes Monte Carlo spin dynamics) was repeated with 9TI4 (Nguyen et al., 2026, 2.66 Å), an independently solved, higher-resolution complex I structure, in place of the manuscript's reference structure 5XTD (Guo et al., 2017, 3.70 Å), with all physical parameters and rate constants unchanged. (A) Normalised O₂ steric accessibility, A(r), from 5XTD (grey) and 9TI4 (blue); dashed lines, previous version with FMN atoms omitted. (B) Normalised composite hotspot score, H(r); dotted lines mark each structure's top-decile mean. (C) Distribution of FMN–O₂ distances within the top-10% hotspot ensemble for each structure, with kernel density estimates and means (dashed). With FMN-overlapping positions excluded, the top-decile means are 5.80 Å for 5XTD (2.5th–97.5th percentiles, 5.56–6.04 Å) and 7.86 Å for 9TI4 (7.74–7.98 Å), so the location of the score maximum depends on the structure. The close agreement reported in the previous version (3.28 and 3.25 Å; grid peaks both 3.25 Å) arose because both calculations counted positions overlapping FMN as accessible.

**Supplementary Note 2: Confirmatory 1,300,000-candidate re-analysis and high-precision |ΔY(B)| plateau check (12 Aug 2026)**

Two follow-up analyses were performed to test explanations raised for the broadened hotspot spread reported in the extended 2.5–20 Å domain (Supplementary Note 1, Table 1). First, the structural accessibility ensemble was regenerated using 1,300,000 candidate O_2_ configurations (versus the original 120,000), using the identical element-specific van der Waals steric-filter procedure against the deposited 5XTD coordinates (with FMN atoms included in the corrected analysis; the previous version omitted them), over the same 2.5–20 Å domain. Second, |ΔY(B)|(r) was recalculated directly with the re-implemented spin model (corrected dipolar sign) — independently of the composite hotspot score and its accessibility weighting — at a 10-fold higher Schulten-Wolynes Monte-Carlo sample count (N=1,500 versus the production N=200–400) with common random numbers fixed between the low-field and high-field draws at every distance, to determine whether the previously reported |ΔY(B)| peak locations (4.75 Å / 7.85 Å; main text, Fig. S2) reflect genuine spatial structure or Monte Carlo sampling noise combined with the 9 Å domain truncation used in that calculation.

With FMN atoms included in the steric test, the 1,300,000-candidate ensemble had an accessible fraction of 7.6% over 2.5–20 Å for 5XTD (7.5% for 9TI4; 8.4% when FMN atoms were omitted, as in the previous version, which reported 8.3%). It produced no near-contact score maximum (Supplementary Note 1; main text, Table 1): with exchange alone, the score formed a broad plateau beyond about 5 Å with its highest value at the domain boundary, and with dipolar coupling the maximum lay at 6.05 Å (5XTD) or 7.85 Å (9TI4), with broad top-decile distributions (SD 1.43 and 1.82 Å). An 11-fold increase in candidate count did not narrow the extended-range spread, which therefore does not result from limited sampling density.

The high-precision, common-random-number recalculation resolved the actual cause (Fig. S4). In the re-implemented spin model with the corrected dipolar sign, exchange-only |ΔY(B)| rises steeply from near contact for k_STD_ ≤ 3 × 10^7^ s^−1^ (where exchange coupling dominates and suppresses singlet–triplet interconversion) and reaches a plateau by about 5 Å that remains constant to within Monte Carlo precision out to 20 Å. With the collinear dipolar term, the plateau is approached further out (about 8–10 Å) because D(r) decays more slowly than J(r), and a broad maximum appears between about 6 and 13 Å, moving to shorter separation as k_STD_ increases, followed by a slow decline; a narrow feature at 2.75 Å lies inside the region excluded by FMN. In neither model is there a narrow maximum within the accessible range. The previously reported 4.75 Å/7.85 Å “peak” locations (main text; Supplementary Fig. S2) are therefore attributable to the combination of (i) the original calculation's 2.5–9.0 Å domain truncation, which stopped sampling near where the plateau is first reached, and (ii) Monte Carlo sampling noise (N=200–400) selecting a particular near-plateau bin as the nominal maximum — not to a genuine, spatially localised field-sensitivity optimum.

To assess whether any residual population-weighted signal remains in the 4.75–7.85 Å window once structural accessibility is accounted for, |ΔY(B)|(r) was also weighted pointwise by the normalised accessibility profile, A(r), from the 1,300,000-candidate ensemble (population-weighted |ΔY(B)| = |ΔY(B)|·A(r); the same accessibility term used in the composite hotspot score, H(r), in the main text). With FMN-overlapping positions excluded, A(r) is negligible below about 3.5 Å and rises to its maximum near 5.8 Å (5XTD) or 7.8 Å (9TI4), so the population-weighted curve is no longer dominated by the near-contact region. Across the k_STD_ values tested, 19% (5XTD) and 13% (9TI4) of the total population-weighted |ΔY(B)| mass over 2.5–20 Å fell within the 4.75–7.85 Å window with exchange alone (4–22% with exchange plus dipolar coupling), and at most 0.2% fell within 2.77–3.64 Å, because almost no accessible positions remain there. The previous version, in which FMN-overlapping positions were counted as accessible, reported 21–25% and 9–12% (exchange-only) or 0.1–5% (exchange + dipolar), respectively. The 4.75–7.85 Å region therefore carries no disproportionate population-weighted sensitivity relative to its width.

Taken together, these results do not support reframing 4.75–7.85 Å as a distinct “magnetically responsive regime” separate from a near-contact “formation regime”: the region in question is not a spatially confined feature at all, but the point at which exchange (and, when included, dipolar) suppression of |ΔY(B)| has decayed away, after which the raw magnetic-field response is essentially featureless out to the edge of the sampled domain. After correction of the steric test, the composite score has no near-contact maximum. Its maximum lies where accessibility rises and the raw field response reaches its plateau (5.5–6.0 Å in 5XTD and 7.7–8.0 Å in 9TI4 over 2.5–9.0 Å), and over 2.5–20 Å the exchange-only score has no localized maximum. The near-contact hotspot reported in the previous version arose because positions overlapping FMN were counted as accessible.

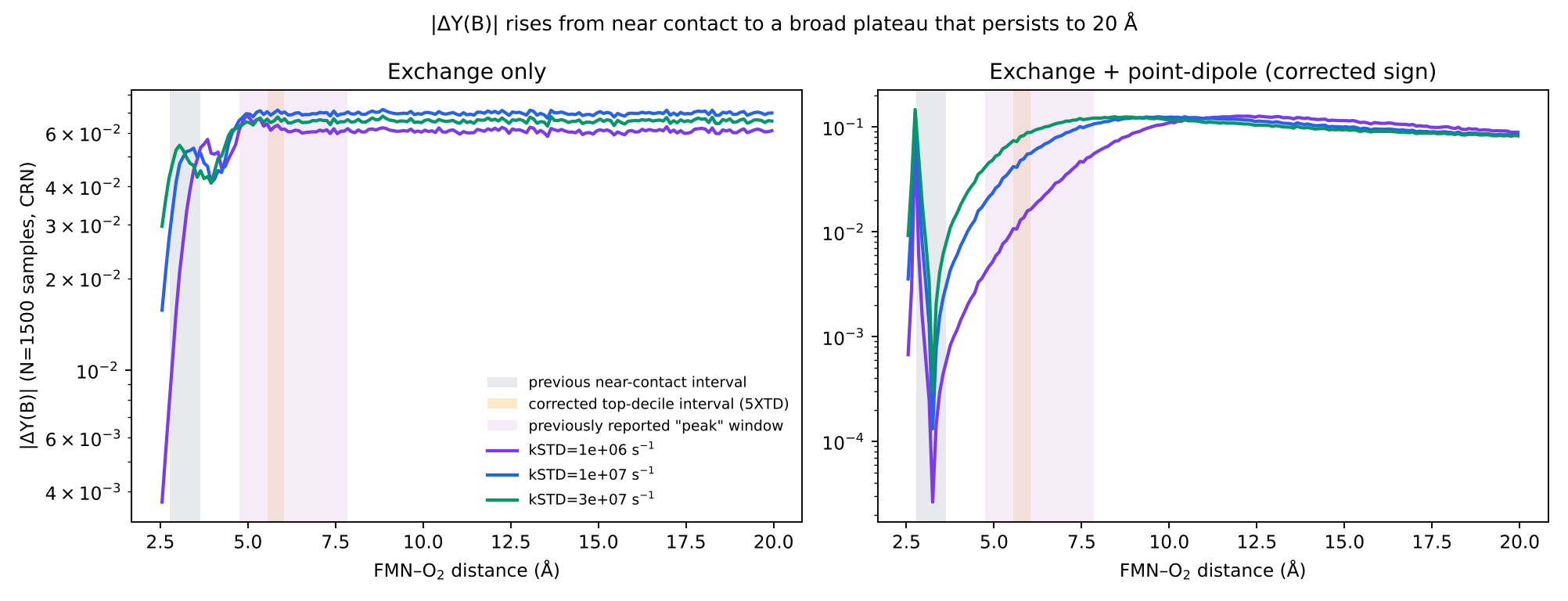

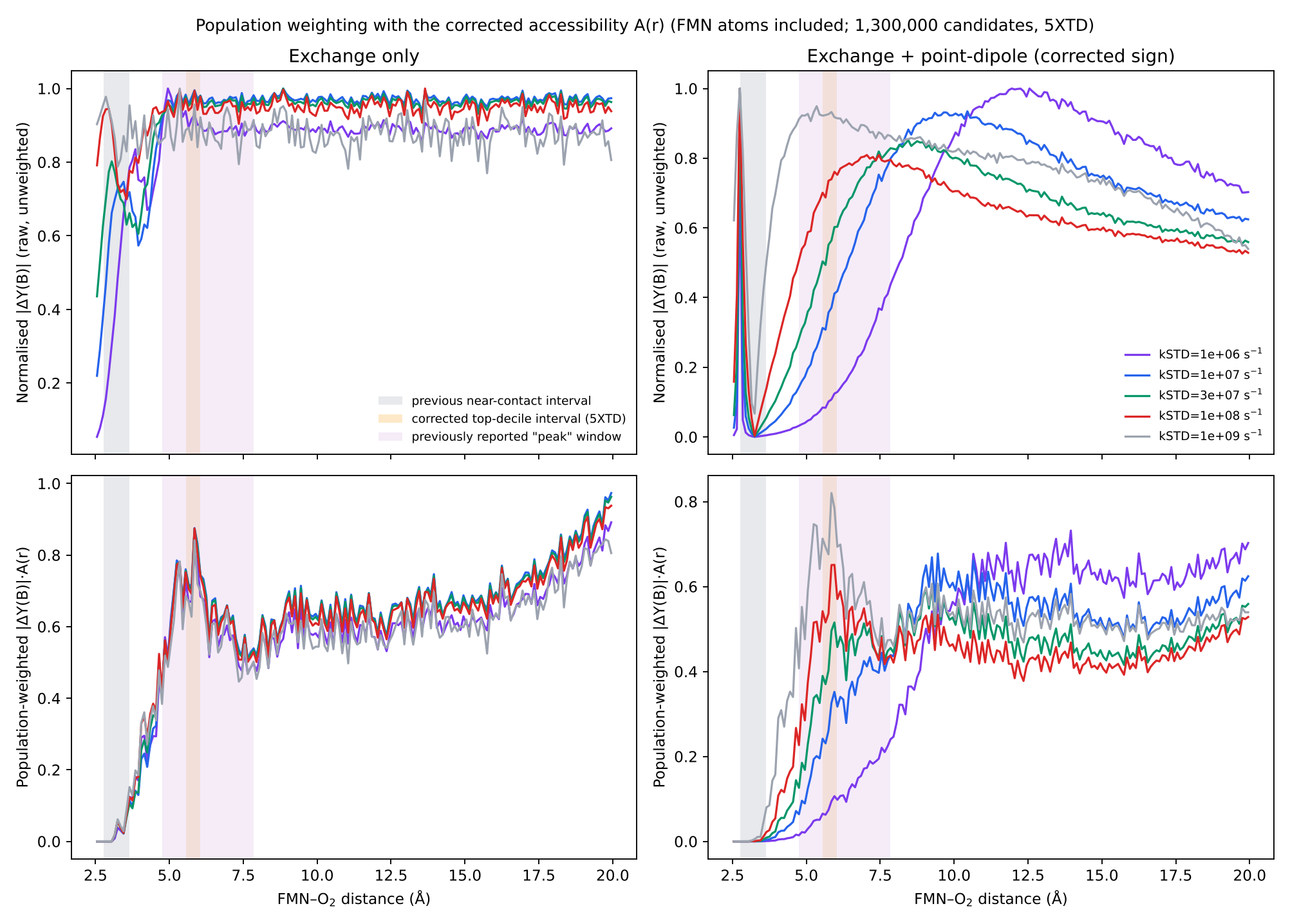

**Figure S4.** Higher-precision field response across 2.5–20 Å (re-implemented spin model; 1,500 Schulten–Wolynes samples per distance with common random numbers). Top: raw |ΔY(B)| for k_STD_ = 10^6^, 10^7^ and 3 × 10^7^ s^−1^, using exchange alone or exchange with a collinear point-dipole term (corrected sign). Bottom: normalised raw |ΔY(B)| (upper row) and accessibility-weighted response |ΔY(B)|·A(r) (lower row) for k_STD_ = 10^6^–10^9^ s^−1^, using the corrected accessibility of 5XTD (FMN atoms included; 1,300,000 candidates). Grey band, near-contact interval of the previous version (2.77–3.64 Å); orange band, corrected 5XTD top-decile interval (5.56–6.04 Å); purple band, 4.75–7.85 Å range of the largest sampled bins in Fig. S2. The raw responses approach broad plateaus without a narrow maximum in the accessible range; the narrow feature at 2.75 Å in the dipolar model lies inside the region excluded by FMN, and accessibility weighting removes all contribution below about 3.5 Å.

**Supplementary Note 3: Correction of the steric test and comparison of two complex I structures**

In the previous version of this analysis, the steric test compared candidate O_2_ centroids with protein heavy atoms but omitted the atoms of the FMN cofactor, from whose isoalloxazine centre the separation r is measured. Of the positions this test accepted within 3.3 Å of the ring centre, 99.4–100% overlapped FMN atoms in 5XTD and 9TI4 (median distance to the nearest FMN atom, 1.2–1.35 Å, compared with the 3.04–3.32 Å required by the van der Waals criterion). With FMN atoms included, ≤0.2% (5XTD) and 0% (9TI4) of candidates within 3.3 Å were accessible, and the accessibility maxima moved to about 5.8 and 7.8 Å, respectively (Fig. S5A). The composite-score top-decile means became 5.80 Å for 5XTD (2.5th–97.5th percentiles, 5.56–6.04 Å) and 7.86 Å for 9TI4 (7.74–7.98 Å) (Fig. S5B), using the spin-response profiles of the original calculation; recombining those profiles with the uncorrected accessibility reproduced the previous results (top-decile mean 3.24 Å for 5XTD; previously 3.28 Å). An independent calculation on a regular 0.1 Å grid with brute-force distance evaluation and a separately written coordinate parser reproduced these results (accessible fractions 5.98% versus 5.96% for 5XTD). Within each structure, the location was insensitive to the O_2_ probe radius (1.40–1.60 Å), FMN radius scaling (0.8–1.0), inclusion of hydrogen atoms and the van der Waals radius set (Fig. S5C), and, in a re-implementation of the spin model, to the dephasing model, flavin hyperfine parameter set (FMN•⁻, 1.233 mT; FMNH•, 1.108 mT) and response-metric definitions (Fig. S5D). The difference between the two structures reflects the modelled pocket: Asn116 and Asn244 lie 3.3 and 3.5 Å from the isoalloxazine centre in 9TI4, compared with 4.5 and 7.4 Å in 5XTD. In the re-implemented spin models, the grid position of the single highest score varied by up to 0.5 Å with sampling seed, bin width and smoothing, so top-decile statistics are reported as the primary measure.

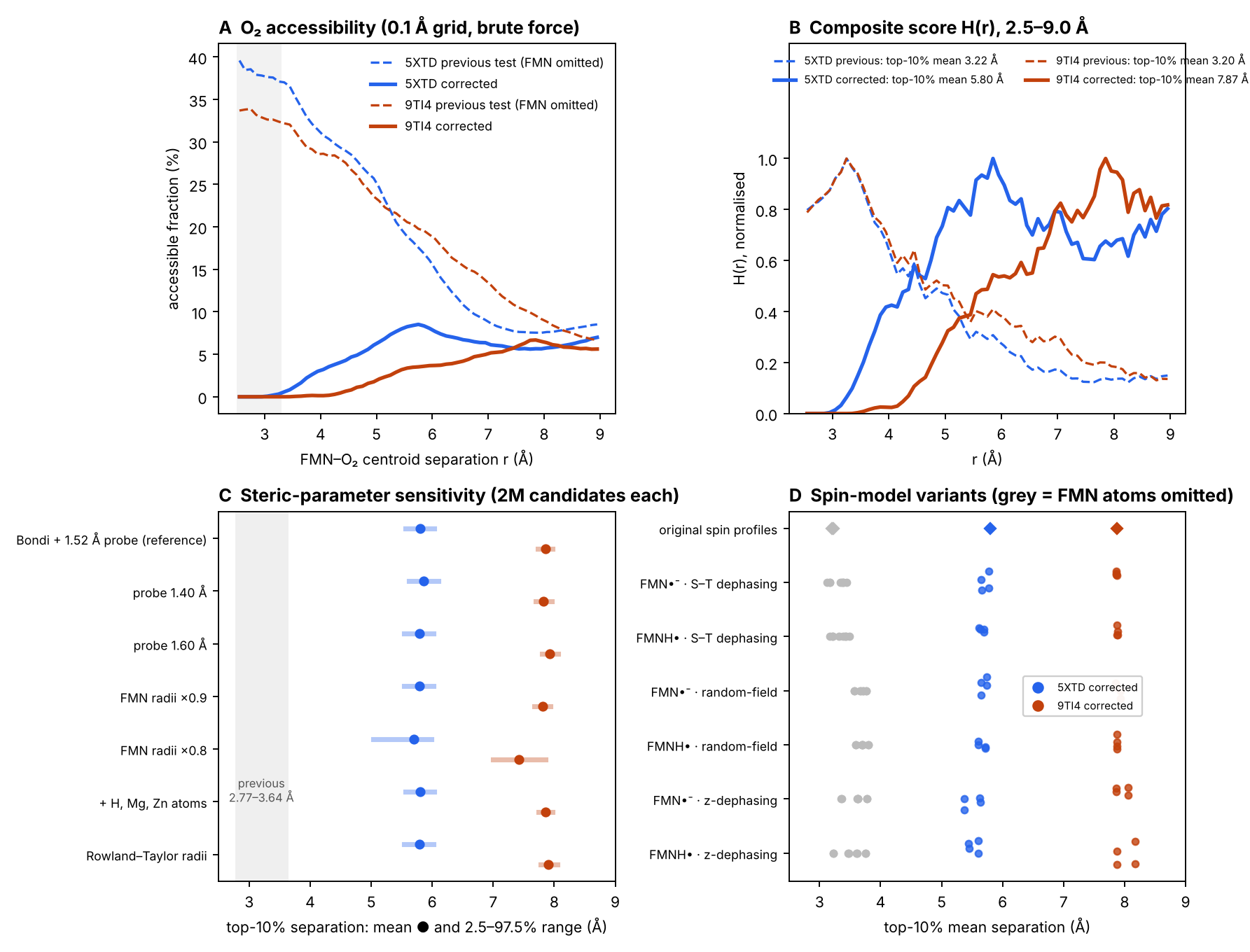

**Figure S5.** Correction of the steric test in two complex I structures. (A) Accessible fraction of candidate O_2_ centroids on a regular 0.1 Å grid, calculated by brute-force distance evaluation, with FMN atoms omitted (dashed; previous version) or included (solid). (B) Composite score H(r) over 2.5–9.0 Å, using the original spin-response profiles, for 5XTD (blue) and 9TI4 (orange), with FMN atoms omitted (dashed) or included (solid). (C) Top-decile mean separations (dots) and 2.5th–97.5th percentile ranges (bars) for steric-parameter variants, each with 2,000,000 candidates. Grey band, near-contact interval reported in the previous version. (D) Top-decile mean separations with the original spin-response profiles (diamonds) and for re-implemented spin-model variants (three dephasing models × two flavin hyperfine sets × four response-metric definitions); grey, FMN atoms omitted.
